# RAN translation of the expanded CAG repeats in the SCA3 disease context

**DOI:** 10.1101/2020.05.07.082354

**Authors:** Magdalena Jazurek-Ciesiolka, Adam Ciesiolka, Alicja A. Komur, Martyna O. Urbanek-Trzeciak, Agnieszka Fiszer

## Abstract

Spinocerebellar ataxia type 3 (SCA3) is a progressive neurodegenerative disorder caused by a CAG repeat expansion in the *ATXN3* gene encoding the ataxin-3 protein. Despite extensive research the exact pathogenic mechanisms of SCA3 are still not understood in depth. In the present study, to gain insight into the toxicity induced by the expanded CAG repeats in SCA3, we comprehensively investigated repeat-associated non-ATG (RAN) translation in various cellular models expressing translated or non-canonically translated *ATXN3* sequences with an increasing number of CAG repeats. We demonstrate that two SCA3 RAN proteins, polyglutamine (polyQ) and polyalanine (polyA), are found only in the case of CAG repeats of pathogenic length. Despite having distinct cellular localization, RAN polyQ and RAN polyA proteins are very often coexpressed in the same cell, impairing nuclear integrity and inducing apoptosis. We provide for the first time mechanistic insights into SCA3 RAN translation indicating that *ATXN3* sequences surrounding the repeat region have an impact on SCA3 RAN translation initiation and efficiency. We revealed that RAN translation of polyQ proteins starts at non-cognate codons upstream of the CAG repeats, whereas RAN polyA proteins are likely translated within repeats. Furthermore, integrated stress response activation enhances SCA3 RAN translation. We suggest that RAN translation in SCA3 is a common event substantially contributing to SCA3 pathogenesis and that the *ATXN3* sequence context plays an important role in triggering this unconventional translation.

## INTRODUCTION

Spinocerebellar ataxia type 3 (SCA3) is the most common form of dominantly inherited ataxia in the world and one of nine polyglutamine (polyQ) neurodegenerative diseases. This progressive and fatal disorder is primarily characterized by neuronal dysfunction and degeneration of the cerebellum and functionally related brain regions. SCA3 is caused by an expanded stretch of CAG repeats in exon 10 of the *ATXN3* gene, which encodes the deubiquitinating enzyme ataxin-3. These repeat tracts range in *ATXN3* from 12 to 44 triplets in healthy individuals and from 56 to 87 in SCA3 patients [1–4]. Until recently, it was believed that the expanded CAG repeats in SCA3 exerted their pathogenic effects at only the protein level. The presence of an abnormally long tract of glutamine in the C-terminal region of ataxin-3 leads to a toxic gain-of-function of the protein, which manifests mainly as characteristic neuronal cytoplasmic and nuclear aggregates [1–4]. However, there is also mounting evidence to suggest that mutant transcripts contribute to the pathogenesis of SCA3 *via* an RNA gain-of-function mechanism [1, 5, 6]. RNA toxicity in this polyQ disease can be induced by aberrant splicing of MBNL1-dependent genes due to the sequestration of this alternative splicing factor by mRNAs containing expanded CAG repeats [7, 8]. Moreover, an abnormal interaction between mutant *ATXN3* transcripts and nucleolin, resulting in the induction of nucleolar stress, was reported in a SCA3 model [9].

The pathogenesis of SCA3 is expected to be more complex, as additional mechanisms are being discovered in other repeat expansion diseases that may also function in SCA3. One such example is repeat-associated non-ATG (RAN) translation, which was described for repeat tracts of CAG, CUG, CGG, CCG, GGGGCC, and CCCGG and, recently, for CCUG, CAGG and UGGAA units [10–19]. These expanded repeat tracts, located mainly in the non-coding regions of specific genes, can trigger non-canonical translation, i.e., in the absence of an AUG initiation codon induce the synthesis of various sets of potentially toxic proteins composed of homopolymeric, dipeptide, tetrapeptide or even pentapeptide repeated tracts of amino acids. RAN translation can be initiated in multiple reading frames in both sense and antisense directions, leading to the expression of distinct proteins [10–19]. The precise mechanism of RAN translation initiation is still poorly understood; however, it has been shown that the repeat tract length and RNA structure formed by repeats play crucial roles [20–23]. More recently, it was revealed that RAN translation of expanded CGG and GGGGCC repeats might utilize a cap-dependent and/or cap-independent ribosomal scanning mechanism for the initiation at near-cognate codons upstream of the repeats [24–30]. Furthermore, among the potential regulators of RAN translation process identified i.a. by genetic screens are translation initiation factors, DDX3X helicase and ribosomal protein S25 [31–35]. Additionally, the cellular stress response promotes this non-canonical mode of protein synthesis from CGG and GGGGCC repeats [26, 27, 29, 36]. A large body of evidence shows the high relevance of RAN proteins for neurodegeneration; these proteins form aggregates that accumulate in both the affected tissues of patients and animal models of SCA8, myotonic dystrophy type 1 (DM1), *c9orf72* amyotrophic lateral sclerosis/frontotemporal dementia (c9ALS/FTD), fragile X tremor ataxia syndrome (FXTAS), Huntington’s disease (HD), DM2, SCA31 and Fuchs’ endothelial corneal dystrophy (FECD) [37–39]. Depending on the type of repeats and the experimental model used, RAN protein toxicity is attributed to the impaired function of the ubiquitin proteasome system, aberrant nucleocytoplasmic transport or the induction of nucleolar and endoplasmic reticulum stress and usually results in enhanced cell death [37–41].

RAN translation is mainly described for diseases in which repeat expansion is located in non-coding regions of the associated genes. In polyQ disorders, in which repeat regions are present in open reading frames, RAN translation has not been extensively studied. There has been one report describing the importance of RAN translation in HD patients, and the initial results of another study suggested that this phenomenon might also occur in SCA2 [15, 42].

In the present study, we show that RAN translation is initiated at CAG repeats located in the sequence context of the *ATXN3* gene and might represent an additional toxic pathway involved in the pathology of SCA3. To show this, we generated multiple translated or non-canonically translated *ATXN3* constructs with different numbers of CAG repeats and various lengths of surrounding sequences and utilized them to analyze RAN translation in transfected cells. We demonstrate that SCA3 RAN translation occurs exclusively in cells expressing constructs containing pathogenic CAG repeats, and the efficiency of this process greatly depends on the *ATXN3* sequence flanking the repeat region. Interestingly, the resulting RAN proteins, i.e., polyQ and polyalanine (polyA) products exhibit different cytoplasmic and nuclear localization; however they very often coexist within the same cell. Importantly, in cells expressing these unusual proteins, the nuclear envelope structure is disturbed. Proteomic analysis revealed that RAN translation in the glutamine frame is initiated from non-cognate codons upstream of the repeat region. We also found that various cellular stress conditions upregulate CAG RAN translation through eIF2α phosphorylation.

## MATERIALS AND METHODS

### Generation of the *ATXN3* constructs

Full-length human *ATXN3* cDNA was previously cloned from total mRNA isolated from SCA3 patient-derived fibroblasts (GM06153, obtained from NIGMS Human Genetic Cell Repository at the Coriell Institute for Medical Research) [43]. The cloning strategy used for the constructs expressing translated or untranslated *ATXN3* cDNA containing different lengths of CAG tracts was previously described [8, 44]. These constructs were prepared in a pCDH-EF1α-MCS-BGH-PGK-GFP-T2A-Puro vector (System Biosciences). To monitor the expression of every possible RAN protein, these constructs were modified by cloning into an Eco0109I site, which is located downstream of the repeat region, a DNA oligo encoding three different protein tags (HA-tag for glutamine, Myc-tag for alanine and His-tag for serine reading frames). In the case of constructs containing shortened 5’ and 3’ flanking sequences, first, a sequence containing the same three tags and two stop codons in each frame was cloned into a pcDNA3.1 (Invitrogen) expression vector using EcoRI and Eco0109I restriction sites. Second, inserts containing fragments of the *ATXN3* sequence with various lengths of CAG repeats were amplified using the full-length cDNA of *ATXN3* as template constructs and specific primers that also introduced BamHI and EcoRI restriction sites at the 5’ and 3’ ends, respectively. Additionally, a forward primer was designed to add two stop codons before the 5’ flanking sequence in each frame. PCR products were cut with BamHI and EcoRI and cloned into a pcDNA3.1 vector. Mutation of the near-cognate codon AAG to CAG was performed using a QuikChange II XL Site-Directed Mutagenesis Kit (Agilent). All generated constructs were verified by Sanger sequencing.

### Cell culture

HEK293T and HeLa cells (ATCC) were grown in DMEM (Gibco) supplemented with 8% FBS (Biowest), 2 mM L-glutamine (Gibco) and 100 U/ml penicillin/100 μg/ml streptomycin (Gibco). Human neuroblastoma SH-SY5Y cells (ATCC) were grown in DMEM/F12 medium (Gibco) supplemented with 10% FBS, 100 U/ml penicillin/100 μg/ml streptomycin (Gibco), and 1x non-essential amino acids (Sigma-Aldrich). The cells were maintained under standard conditions of temperature (37°C), humidity (95%) and carbon dioxide (5%). For longer storage periods, the cells were frozen in freezing medium and stored in liquid nitrogen.

### Plasmid transfection and drug treatments

Routine cell transfection of HEK293T and HeLa cells with plasmids was conducted using Lipofectamine 2000 and Lipofectamine LTX (Life Technologies), respectively, according to the manufacturers’ instructions. Prior to transfection, cells were plated at the required confluence on 12-well plates and harvested 48 h after transfection. For stress induction, HEK293T cells were transfected with the generated constructs for 20 h, followed by 8 h of treatment with 1 μM thapsigargin (Enzo Life Technologies) or 50 μM sodium arsenite (Sigma-Aldrich). SH-SY5Y cells were electroporated with a NeonTM Transfection System (Invitrogen) in 100-μl tips using the following parameters: 1.100 V, 50 ms, and 1 pulse.

### Western blotting

Routinely, cells were sonicated in lysis buffer containing a 60 mM TRIS-base, 2% SDS, 10% sucrose, 2 mM PMSF and phosphatase inhibitor cocktail (PhosSTOP; Roche). For the analysis of RAN translation after cellular stress induction, the cells were lysed in RIPA buffer (Sigma-Aldrich) containing protease inhibitor cocktail (cOmplete, EDTA-free; Roche) and phosphatase inhibitor cocktail. The protein concentration was estimated using a NanoDrop spectrophotometer. A total of 50 μg of protein was diluted in sample buffer containing 2-mercaptoethanol, denatured at 95°C for 5 min, and separated on a 12% SDS-PAGE gel. After electrophoresis, the proteins were transferred onto a 0.1 μm nitrocellulose membrane (Amersham), blocked with 5% non-fat dry milk in TBS-0.1% Tween-20 for 1 h and incubated overnight with mouse anti-ataxin-3 (1:1000; Millipore), mouse anti-HA (1:1000; Sigma-Aldrich), rabbit anti-c-Myc (1:1000, Cell Signaling Technology, CST), rabbit anti-His (1:1000, CST), rabbit anti-phospho-eIF2α (1:1000, CST), mouse anti-eIF2α (1:1000, CST) or mouse anti-GAPDH (1:20000, Millipore) antibodies. Next, the membranes were incubated with secondary anti-mouse or anti-rabbit (Jackson ImmunoResearch) peroxidase-conjugated antibodies for 1 h, at room temperature (RT). The immunoreaction was detected using the WesternBright Quantum HRP Substrate (Advansta) and G:BOX chemiluminescent camera (Syngene). Band intensity was measured using ImageJ software.

### Mass spectrometry analysis

To immunoprecipitate AUG-initiated and RAN proteins (polyQ and polyA), HEK293T cells transfected with AUG-78/32-117CAG, 78/32-115CAG or 33/32-119CAG constructs were lysed after 48 hours in RIPA buffer containing protease and phosphatase inhibitor cocktails. HA-tagged polyQ and Myc-tagged polyA proteins were enriched using Pierce™ Anti-HA Agarose (Thermo Fisher Scientific) and Myc-Trap®_A Kit (Chromotek) respectively, according to the manufacturer’s instructions. After washing steps the agarose beads bound with polyQ and polyA proteins were further processed for LC-MS/MS analysis. Mass spectrometry experiments were performed and analyzed at the Mass Spectrometry Laboratory at the Institute of Biochemistry and Biophysics Polish Academy of Sciences. Agarose beads suspended in 40 μl of immunoprecipitation washing solution were filled up to 150 μl with 100 mM ammonium bicarbonate buffer. Cysteine bridges were reduced by 1 h incubation with 5 mM tris(2-carboxyethyl)phosphine (TCEP) at 60°C followed by 30 min alkylation at RT in the dark with 15 mM iodoacetamide (IAA). The samples were mixed thoroughly and divided into equal parts for digestion with various enzymes selected on the basis of the distribution of potential cleavage sites in the regions surrounding the polyQ and polyA sequences. All enzymes were of sequencing grade purity and were purchased from Promega Corporation. PolyQ samples were digested with 0.01 μg/μl of LysC or 0.005 μg/μl of trypsin. PolyA samples were digested with trypsin and elastase in a final concentration of 0.01 μg/μl. After overnight digestion, solutions were acidified with 0.1% trifluoroacetic acid and centrifuged (20000 g, 15 min, 4°C) to remove agarose beads from peptide solution. Next, 20 μl of each sample was analyzed using nanoACQUITY UPLC (Waters) directly coupled to a QExactive mass spectrometer (Thermo Fisher Scientific). Peptides were trapped and washed on C18 column (180 μm × 20 mm, Waters) with 0.1% formic acid (FA) in water as a mobile phase and transferred to a nanoACQUITY BEH C18 column (75 μm × 250 mm, 1.7 μm, Waters) using acetonitrile (ACN) gradient (0 – 35% ACN in 160 min) in the presence of 0.1% FA at a flow rate of 250 nl/min. Acquisition was performed in the data-dependent mode with the maximum of 12 precursors selected for MS2. Full MS scans covering the range of 300–1650 m/z were acquired at a resolution of 70000 with a maximum injection time of 60 ms and an AGC target of 1e6. MS2 scans were acquired at the resolution of 17500 and the AGC target of 5e5. Dynamic exclusion was set to 30 s. Acquired data was pre-processed with Mascot Distiller software (2.6, Matrix Science) and a protein identification was performed with Mascot Search Engine (Matrix Science). The searches were done against a homemade database of all potential polyQ and polyA proteins translated from the transfected constructs and the most frequent contaminations (6555 sequences). To reduce mass errors, the peptide and fragment mass tolerance settings were established separately for individual LC-MS/MS runs after a measured mass recalibration, as described previously [45]. The rest of search parameters were as follows: enzyme, Trypsin or Lys-C; missed cleavages, 2 or 3; fixed modifications, Carbamidomethyl (C); variable modifications, Oxidation (M), Deamidation (NQ), Acetyl (N-term), Gln->pyro-Glu; instrument, HCD. Since there was not enough identifications to compute the FDR, peptides identified for polyQ and polyA sequences, for which ion scores were ≥20, were further analyzed manually.

### Immunofluorescence

HEK293T cells were seeded on 12-well plates with glass coverslips (Marienfeld Superior) coated with poly-L-lysine (Sigma-Aldrich) one day prior to transfection. Forty-eight hours post transfection, the cells were fixed with 4% paraformaldehyde for 30 min at RT, permeabilized with 0.5% Triton-X-100 in PBS for 10 min and blocked in 1% BSA for 1 h at RT. The coverslips were incubated overnight at 4°C with the following primary antibodies diluted in 1% BSA in PBS: anti-HA (1:100), anti-c-Myc (1:100), anti-His (1:100), mouse anti-nuclear pore complex (1:1000, BioLegend), and rabbit anti-cleaved-caspase 3 (1:100, CST). The next day, the coverslips were rinsed with PBS and incubated with anti-mouse and anti-rabbit antibodies labeled with Alexa488 and Alexa594, respectively (1:500, Jackson ImmunoResearch), for 1 h at RT. After rinsing the coverslips were mounted with SlowFade Diamond Antifade Mountant with DAPI (Life Technologies). Images were captured with a Leica SP5 confocal microscope and processed using Leica software.

### Statistical analysis

Statistical significance was assessed by an unpaired *t* test and one-way ANOVA with Bonferroni correction for multiple comparisons. A p-value of <0.05 was considered significant. The data are presented as mean ± standard error of the mean (SEM). The results represent at least three independent biological replicates.

## RESULTS

### RAN translation occurs in the *ATXN3* context

To evaluate whether CAG repeats in the sequence context of *ATXN3* can support RAN translation, we developed a series of constructs in pcDNA3.1 plasmid, without a start codon, containing normal (21 CAG) or expanded repeat numbers (61 and 115 CAG) (Fig. 1a, upper scheme). These plasmids contained 78 bp and 32 bp of the *ATXN3* sequence upstream and downstream of the repeat region, respectively. The length of the 5’ flanking sequence was limited by the many ATG codons within the *ATXN3* sequence upstream of the CAG repeat region that were undesired in the constructs, whereas the length of the 3’ flanking sequence was determined with respect to the first stop codon for the alanine reading frame (Fig. S1a). Thus these specific *ATXN3* sequences are expected to be involved in SCA3 RAN translation in case of repeat tract expansion. The generated constructs also contained upstream and downstream stop codons and C-terminal tags located downstream of the 3’ flanking sequence (Fig. 1a). As a control for AUG-initiated translation in the glutamine reading frame, we developed an additional construct with 117 CAG repeats containing a start codon positioned immediately before the 5’ flanking sequence (Fig. 1a, lower scheme). All these constructs were transfected into HEK293T cells, and the potential RAN translation products were analyzed 48 h later.

**Fig. 1.**
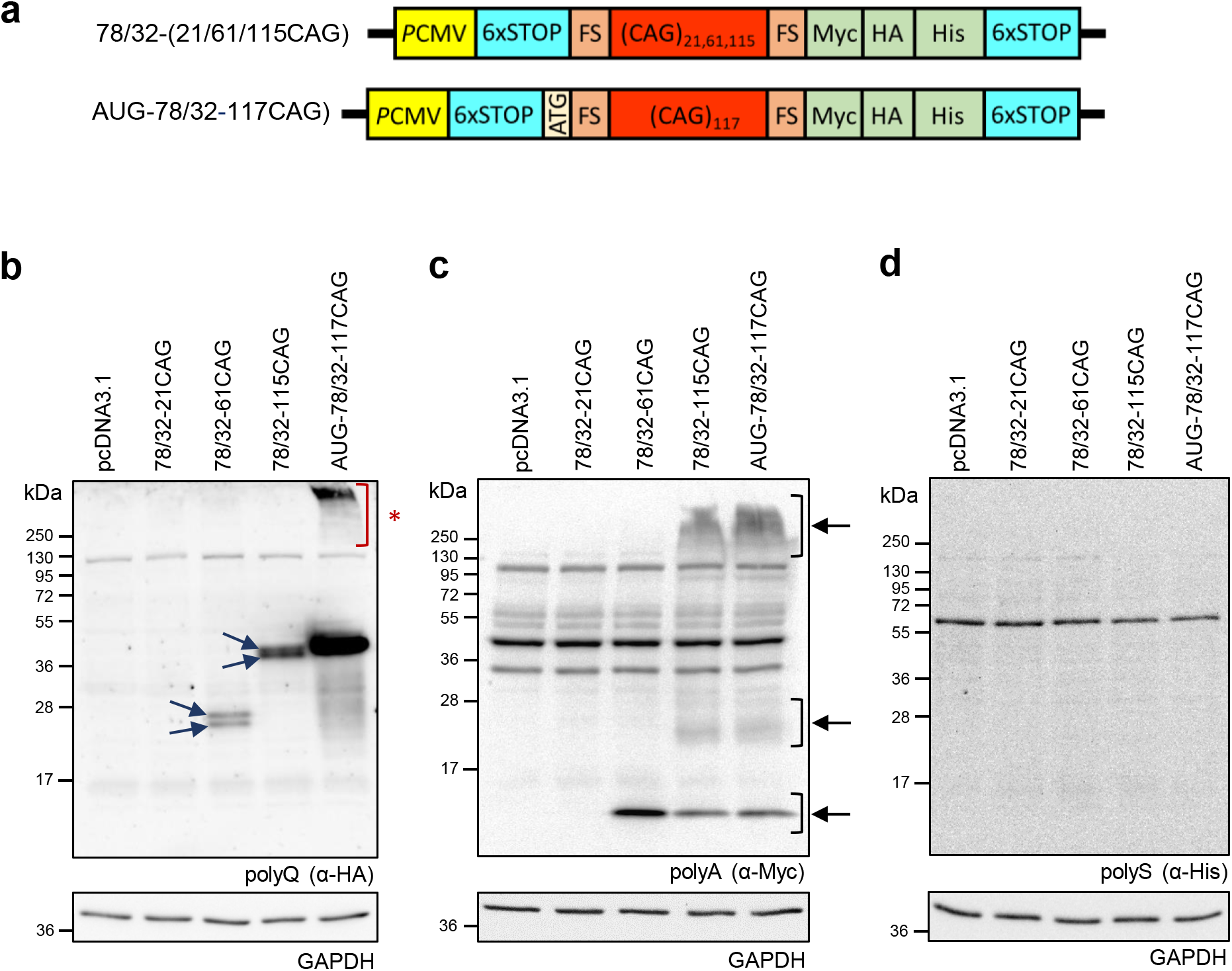
RAN translation of the CAG repeats in the *ATXN3* context. **a** Diagram of the constructs developed to monitor SCA3 RAN translation (upper panel) in reference to canonical translation (lower panel). Both constructs contain 78 bp and 32 bp *ATXN3* flanking sequences (FS) upstream and downstream of repeat region, respectively, and the indicated CAG repeat numbers. Immediately downstream of the *ATXN3*-specific sequence, Myc-, HA- and His-tags were placed in the alanine, glutamine and serine reading frames, respectively. For all the analyses presented in this figure, lysates were obtained from HEK293T cells 48 h after transfection with the indicated constructs. **b** Expression of the AUG-initiated and RAN polyQ proteins was analyzed by western blotting using an anti-HA antibody. RAN translation products are indicated by blue arrows. Aggregated AUG-initiated polyQ proteins are indicated by a red asterisk. **c** Expression of RAN polyA proteins was analyzed by western blotting using an anti-Myc antibody. Specific products are indicated by black arrows. **d** Expression of RAN polyS proteins analyzed by western blotting using an anti-His antibody. No specific signals were detected. For all the western blot analyses, the GAPDH level was used as a loading control

In the absence of an AUG codon, RAN polyQ and RAN polyA proteins were detected for constructs containing expanded CAG repeat tracts (61 and 115 CAG) (Fig. 1b and c). We did not observe any specific signal from potential polyserine (polyS) proteins (Fig. 1d). In the case of the construct with unexpanded repeat tracts (21 CAG), RAN translation did not occur (Fig. 1b–d). RAN polyQ proteins migrated as two bands, for which the molecular weights and amounts increased with increasing CAG repeat tract length (Fig. 1b, Fig. S1b). The level of SCA3 RAN polyQ proteins from the 78/32-115CAG construct was significantly lower than that of the AUG-initiated polyQ proteins (Fig. S1c). Moreover, expression of the AUG-initiated polyQ proteins led to the formation of insoluble aggregates of high molecular weight that also remained in the stacking gel (Fig. 1b). RAN polyA proteins translated from the 61 CAG repeats migrated as one clear band below 17 kDa, whereas for the longer repeats (115 CAG), additional multiple proteins of even higher molecular weight appeared (Fig. 1c). Importantly, RAN polyA proteins were also observed in the cells expressing the AUG-containing construct, which indicates that the presence of the start codon did not inhibit RAN translation in other reading frames (Fig. 1c). The obtained polyA proteins expression pattern was similar to that shown in previous results with respect to other repeat expansion diseases and might suggest that RAN translation in the alanine reading frame initiates at multiple sites within the repeats, leading to the generation of products of diverse molecular weights [10, 11, 15, 24].

### SCA3 RAN polyQ and RAN polyA proteins exhibit different subcellular localization and aggregation properties

To determine the subcellular distribution of SCA3 RAN polyQ and RAN polyA proteins, we performed immunofluorescence analysis. RAN polyQ proteins were distributed throughout the cytoplasm and nucleus, whereas RAN polyA proteins were exclusively cytoplasmic (Fig. 2a, Fig. S2). Moreover, SCA3 RAN polyQ proteins exhibited aggregation properties; however, the number of aggregates was fewer compared to those formed by the AUG-initiated polyQ proteins and was dependent on the length of the repeat tract (Fig. 2a and b, Fig. S2). Approximately 3% and 22% of the cells contained aggregates formed by RAN polyQ proteins translated from constructs bearing 61 and 115 CAG repeats, respectively (Fig. 2b). We did not detect inclusions formed by RAN polyA proteins; however, we cannot exclude the possibility that these proteins exist as soluble oligomers (Fig. 2a, Fig. S2) [46–48]. Importantly, SCA3 RAN polyQ and RAN polyA proteins were very often coexpressed in the same cells, which might suggest that their translation was triggered by the same mechanism (Fig. 2a, Fig. S2). Moreover, RAN polyA proteins were also detected in cells containing AUG-initiated polyQ proteins (Fig. 2a, Fig. S2). As expected, we did not find RAN translation products in the cells expressing the nonpathological length of CAG repeats (21 CAG) (Fig. S2). Moreover, as revealed previously by western blot analysis, we did not detect any polyS proteins (data not shown).

**Fig. 2.**
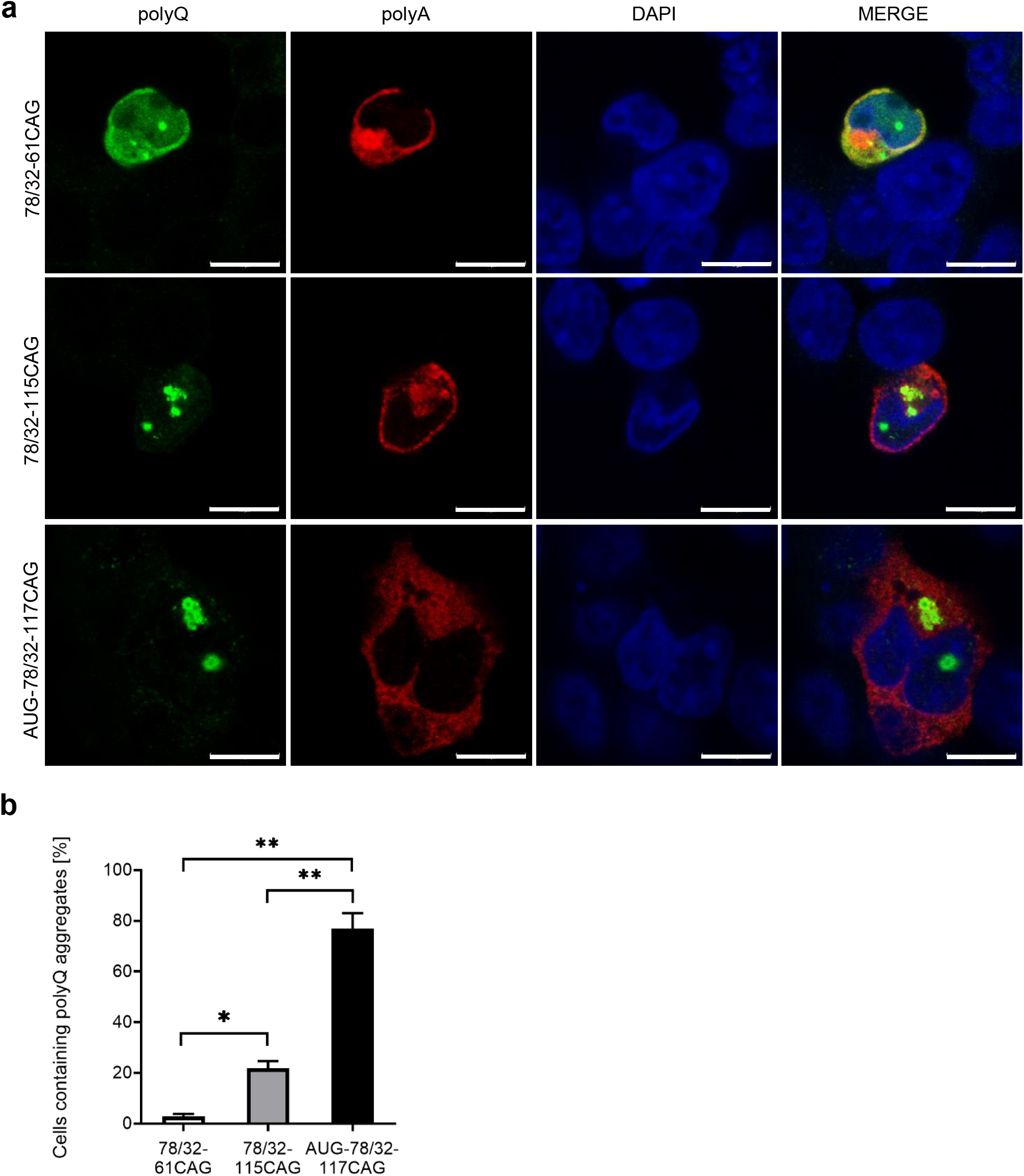
SCA3 RAN polyQ proteins form aggregates. **a** Representative images of immunofluorescence staining for AUG-initiated and RAN polyQ proteins (green signal, anti-HA antibody) and RAN polyA proteins (red signal, anti-Myc antibody) in the HEK293T cells 48 h after transfection with the indicated constructs. DAPI (blue signal) was used to stain nuclei. Scale bar=10 μm. **b** Quantitation results of the cells containing polyQ aggregates; 100% represents cells expressing AUG-initiated or RAN polyQ proteins. The graph bars represent the mean value ± SEM from 4 biological replicates. One-way ANOVA with Bonferroni’s multiple comparisons, *p<0.05, **p<0.0001

### SCA3 RAN proteins affect nuclear integrity and cause apoptosis

Many reports show that RAN proteins are toxic to cells [37–41]. During a microscopic analysis, we observed that, in the cells expressing SCA3 RAN proteins, the shape of the nucleus was disturbed (Fig. 2, Fig. S2). It had been previously shown that perinuclear aggregates formed by polyQ-expanded proteins induce focal distortion of the nuclear envelope [49]. More recently, it was reported that mutant huntingtin disrupts nuclear architecture, which results in aberrant nucleocytoplasmic transport [50, 51]. To analyze the nuclear membrane integrity in the cells expressing SCA3 RAN proteins, immunofluorescence staining of nuclear pore complex (NPC) proteins was performed. As RAN polyQ and RAN polyA proteins frequently coexist in the same cell (Fig. 2a, Fig. S2), we studied the correlation of the nuclear morphology with the RAN polyA proteins only. We found that the detection of RAN polyA proteins was associated with altered NPC localization, which resulted in defects in nuclear envelope assembly (Fig. 3). In contrast, in the cells transfected with empty plasmid or expressing CAG repeats of normal length, the nuclear integrity was not disturbed (Fig. 3).

**Fig. 3.**
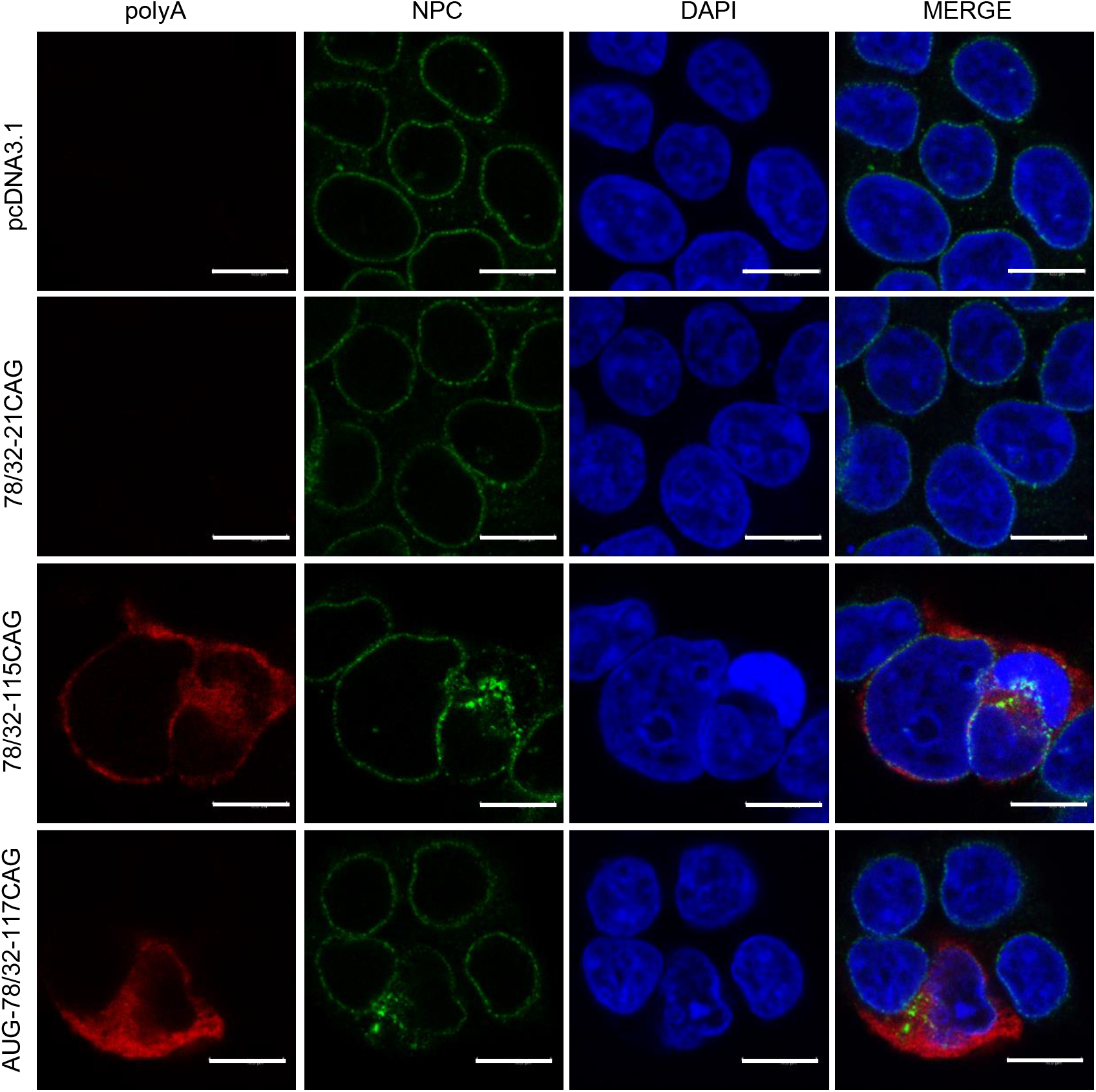
SCA3 RAN polyA proteins affect the integrity of the nuclear membrane. Representative images of immunofluorescence staining for RAN polyA proteins (red signal, anti-Myc antibody), and the nuclear pore complex (NPC, green signal) in the HEK293T cells 48 h after transfection with the indicated constructs. DAPI (blue signal) was used to stain nuclei. Scale bar<10 μm

Disorganization of the nuclear envelope is often connected with the apoptosis process [52]. To address whether SCA3 RAN translation can trigger apoptosis, we additionally analyzed the presence of cleaved caspase-3, a well-known marker of apoptosis induction. As revealed by immunofluorescence, approximately 40% and 8% of the cells expressing AUG-containing and AUG-lacking constructs, respectively, were positive for both cleaved caspase-3 and polyQ proteins (Fig. S3a and b). This finding suggests a substantial impact of non-canonical translation on apoptosis induction, as the cumulative effect observed for the AUG construct was attributed to the presence of both canonically translated and RAN-translated proteins. In conclusion, our data suggest that the expression of SCA3 RAN proteins substantially affects nuclear integrity and promotes cell death.

### SCA3 RAN translation is influenced by the sequences surrounding the expanded CAG repeat region

In the next step, we verified whether RAN translation could also be observed for the full-length *ATXN3* transcript in our experimental setup. We used constructs with various lengths of CAG repeats containing translated (20, 60, 85 and 122 CAG repeats) and non-canonically translated (20, 60, 91 and 126 CAG repeats) full-length *ATXN3* cDNA. Sequences encoding three epitope tags were added immediately downstream of the CAG repeats (Fig. S4a). These constructs were transfected into HEK293T cells, and the level of SCA3 RAN translation was analyzed 48 h after transfection. The cells expressing the translated *ATXN3* constructs produced one strong band corresponding to the overexpressed exogenous ataxin-3 with various lengths of polyQ tracts and additional multiple bands, which may have represented proteins produced from alternative initiation sites or post-translationally modified ataxin-3 (Fig. S4b and c). However, no specific RAN polyA or RAN polyS proteins were detected in cells transfected with these constructs (Fig. S4d and e). In the case of constructs without AUG codons, as expected, the exogenous ataxin-3 expression was abolished (Fig. S4f and g). However, still SCA3 RAN translation was not observed in the cells with any of the three possible frames (Fig. S4g-i).

As we observed a significant difference in the occurrence of RAN translation in the full-length *ATXN3* constructs and their shortened versions, we decided to better recognize the sequence requirements for RAN translation initiation in mutant *ATXN3*. We modified the SCA3 RAN construct 78/32-115CAG by removing 33 nt or 60 nt of the 5’ sequence and 15 nt of the 3’ sequence from the repeats (Fig. S5a). HEK293T cells were transfected with these newly established vectors (45/32-110CAG, 45/17-117CAG, 18/32-119CAG and 18/17-116CAG), and 48 h after transfection, the SCA3 RAN translation was analyzed. Deleting 33 nt of the sequence upstream and/or 15 nt of the sequence downstream of the repeat tract (45/32-10CAG and 45/17-117CAG) did not affect RAN translation in the glutamine frame (Fig. S5b). In contrast, when we truncated the 5’ sequence to 18 nt, the RAN translation in the glutamine frame was abolished (Fig. S5b). In the case of the RAN polyA proteins, we did not observe this high dependency between the occurrence of RAN translation and the length of the 5’ flanking sequence (Fig. S5c). However, we found a significant effect of the 3’ flanking sequence on the RAN translation efficiency in the alanine frame. Shortening of the 3’ flanking sequence to 17 nt while maintaining the length of the 5’ end (45/32-110CAG vs 45/17-117CAG and 18/32-119CAG vs 18/17-116CAG) led to a decrease in the level of RAN polyA proteins (Fig. S5c and d). These results show the profound involvement of the *ATXN3* 5’ flanking sequence in the RAN translation initiation in the glutamine frame and the influence of the 3’ flanking sequence on polyA translation efficiency.

To explain the high sequence dependence of SCA3 RAN translation in the glutamine frame, we analyzed the 5’ flanking sequence for the presence of alternative start codons (Fig. S6a). It had been reported that RAN translation of CGG and GGGGCC repeats is initiated at near-cognate codons located upstream of the repeat tract [11, 24–26, 28–30]. Alternative start codons differ from AUG codon by only one nucleotide and generally are less efficient than standard AUG codon [53, 54]. Two near-cognate AAG codons were found in the glutamine frame in the *ATXN3* sequence in 78/32-115CAG, 45/32-110CAG and 45/17-117CAG constructs (Fig. S6a). In the constructs with the *ATXN3* 5’ flanking sequence truncated to 18 nt (18/32-119CAG and 18/17-116CAG), one of these codons was absent, which suggested the potential involvement of this AAG codon in RAN translation initiation (Fig. S6a). To examine this possibility, in the 45/32-110CAG and 45/17-117CAG constructs, we mutated this AAG codon to CAG codon using site-directed mutagenesis. Western blot analysis of the cells transfected with these mutated constructs did not show AAG codon requirements for the initiation of RAN polyQ proteins (Fig. S6b). Additionally, the introduced mutation did not affect RAN translation of RAN polyA proteins (Fig. S6c)

Thus, to determine the minimal sequence requirements for translation initiation in the glutamine frame, we designed an additional set of constructs (33/32-119CAG, 33/17-118CAG, 24/32-116CAG and 24/17-119CAG), in which the 5’ flanking sequence was shortened to 33 nt and 24 nt, whereas the 3’ flanking sequence was shortened to 17 nt (Fig. 4a). When we truncated the 5’ flanking sequence to 33 nt, we were surprised to observe enhanced RAN translation for the polyQ proteins compared to that of the longer constructs, whereas 3’ shortening did not affect translation (Fig. 4b). Interestingly, the resultant RAN polyQ proteins more often formed aggregates, as determined by western blotting and confocal microscopy (Fig. 4b and e). In contrast, when the 5’ flanking sequence was shortened to 24 nt, RAN translation in the glutamine frame was abolished (Fig. 4b). Similar to previous experiments, truncation of the 3’ flanking sequence to 17 nt resulted in decreased RAN translation efficiency in the alanine frame (Fig. 4c and d).

**Fig. 4.**
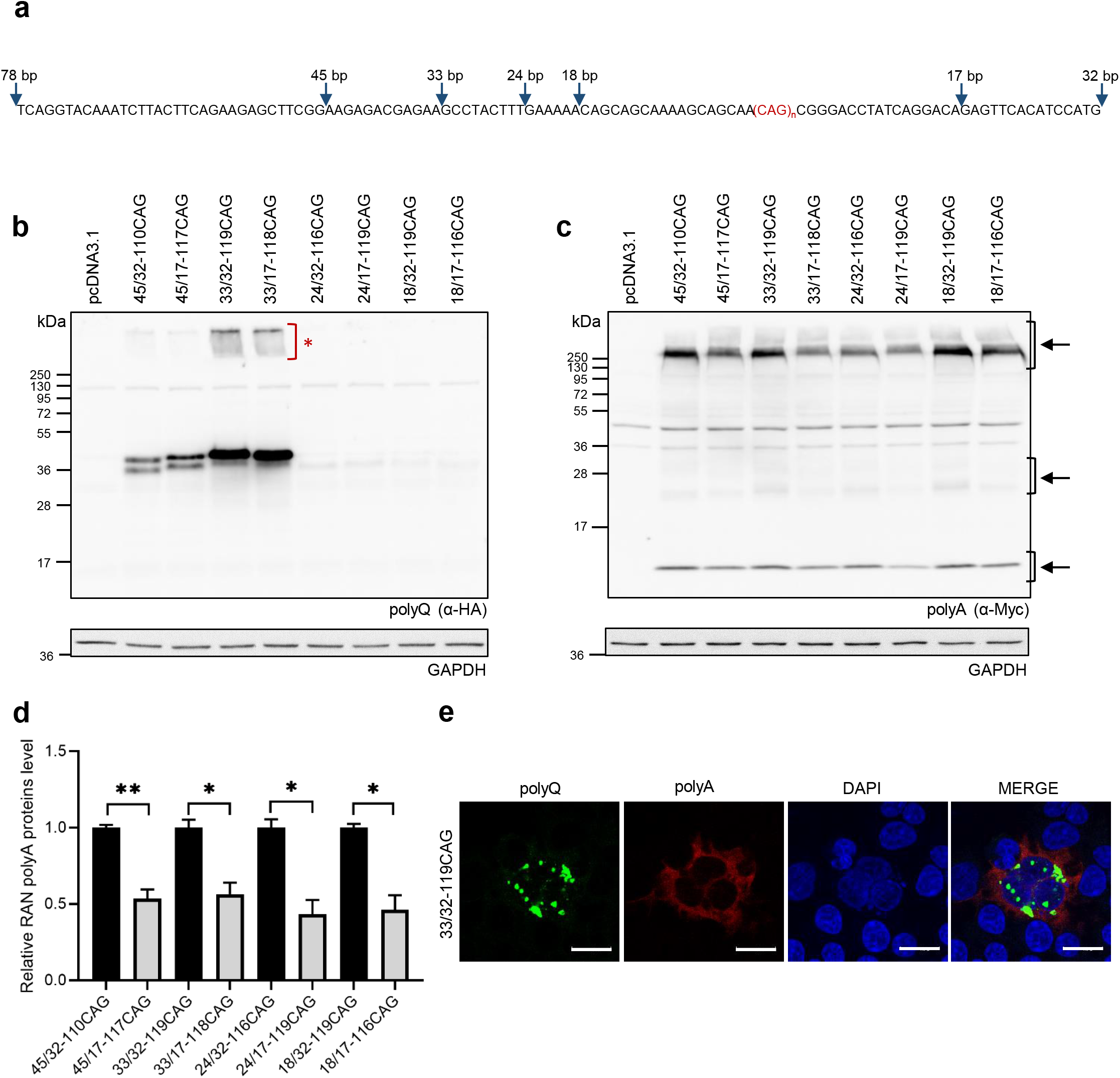
SCA3 RAN translation in the glutamine frame greatly depends on the 5’ flanking sequence. **a** *ATXN3* 5’ and 3’ flanking sequences of the 78/32-115CAG, 45/32-110CAG, 45/17-117CAG, 33/32-119CAG, 33/17-118CAG, 24/32-116CAG, 24/17-119CAG, 18/32-119CAG and 18/17-116CAG constructs. The blue arrows indicate the length of the flanking sequence upstream and downstream of the repeats used in particular RAN translation constructs. For all the analyses presented in this figure, lysates were obtained from HEK293T cells 48 h after their transfection with the indicated constructs. **b** Expression of the RAN polyQ proteins was analyzed by western blotting using an anti-HA antibody. Aggregated RAN polyQ proteins are indicated by the red asterisk. **c** Expression of the RAN polyA proteins was analyzed by western blotting using an anti-Myc antibody. **d** Relative expression of the RAN polyA proteins from the indicated constructs normalized to that of the 45/32-110CAG, 33/32-119CAG, 24/32-116CAG or 18/32-119CAG constructs. Specific RAN polyA products used in the quantitative analysis are indicated by black arrows. The graph bars represent the mean value ± SEM from 4 biological replicates. Two-tailed *t* test, *p<0.005, **p<0.0005. For all the western blot analyses, GAPDH was used as a loading control. **e** Representative images of immunofluorescence staining for RAN polyQ proteins (green signal, anti-HA antibody) and RAN polyA proteins (red signal, anti-Myc antibody) in the HEK293T cells expressing 33/32-119CAG construct. DAPI (blue signal) was used to stain nuclei. Scale bar=20 μm

We also examined the influence of the 5’ flanking sequence on SCA3 RAN translation initiation in the glutamine frame in other cell lines. Transfection of HeLa and SH-SY5Y cells with constructs differing only in the length of the 5’ sequence upstream of the CAG region (78/32-115CAG, 45/32-110CAG, 33/32-119CAG, 24/32-116CAG and 18/32-119CAG) demonstrated that SCA3 RAN translation can proceed in different cellular backgrounds (Fig. S7). However, in contrast to that in the HEK293T cells, RAN translation in the other cell lines, particularly in neuroblastoma cells, was less efficient. This effect can also be explained by differences in the transfection efficiency between the studied cell lines. In the case of the HeLa cells, we were able to detect the signal for AUG-initiated polyQ proteins and RAN polyQ proteins produced from the 33/32-119CAG construct and a very weak signal for RAN polyA proteins expressed from studied RAN constructs (Fig. S7a and 7b). We also observed the same polyQ proteins in SH-SY5Y cells as in the HeLa cells; however, in contrast to the HeLa cells, there was no specific signal for RAN polyA proteins in the SH-SY5Y cells (Fig. S7c and d). Taken together, these data seem to indicate that the 5’ flanking sequence, particularly that located 24 nt upstream of the CAG repeats, plays a significant role in RAN translation initiation in the glutamine frame, and the efficiency of RAN translation might be cell-type dependent.

### SCA3 RAN translation in the glutamine frame is initiated at upstream non-cognate codons

In order to characterize translation initiation sites solely dependent on RAN translation, we used immunoprecipitation of RAN polyQ and RAN polyA proteins from the HEK293T cells transfected with 78/32-115CAG and 33/32-119CAG constructs and analyzed specific peptides by mass spectrometry. Additionally, we immunoprecipitated the AUG-initiated polyQ proteins from the cells transfected with the AUG-78/32-117CAG construct as a control to show the events dependent on canonical translation initiation. To map the initiation sites of RAN and canonical translation products with high confidence, as well as to exclude any bias caused by digestion, we digested immunopurified AUG-initiated and RAN polyQ proteins with two different proteases, trypsin and LysC (Fig. S8a, Fig. S9a, Fig. S10a). Moreover, the RAN polyA proteins were digested with trypsin and with an unspecific protease, i.e., elastase (Fig. S8b, Fig. S9b, Fig. S10b). These proteolytic digestion selections enabled us to identify many unique peptides from both polyQ and RAN polyA proteins, many of which originated from regions downstream of the CAG repeats (Fig. 5, Fig. S8, Fig. S9, Fig. S10). Moreover, the analysis of our LC-MS/MS data compared with that of the human proteome (UniProtKB Release 2020_01) resulted in no unique peptides that could have originated from the proteolytic digestion of the endogenous, full-length ATXN3, confirming that all the analyzed peptides originated from immunoprecipitated AUG-initiated polyQ, RAN polyQ and polyA proteins.

**Fig. 5.**
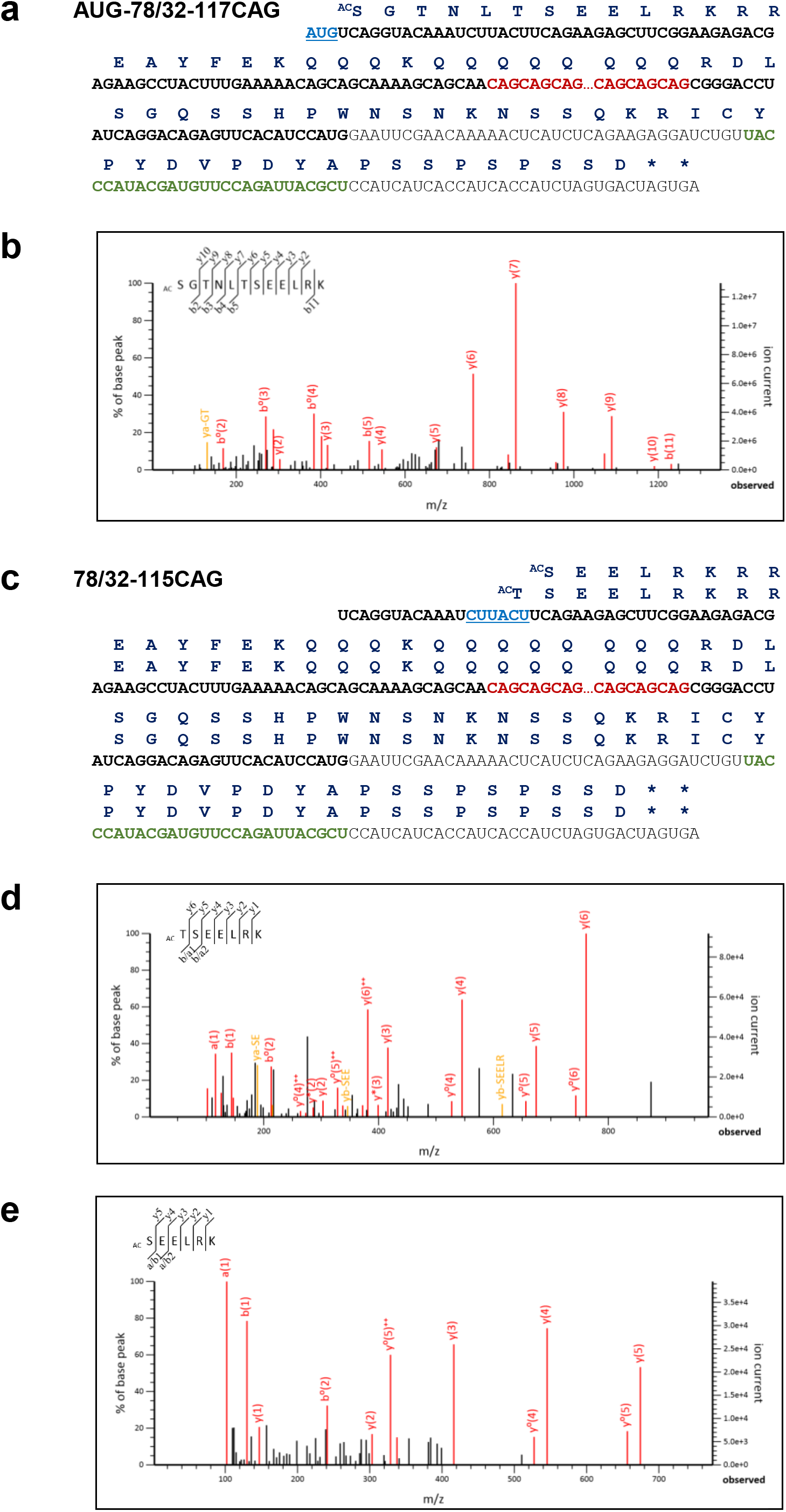
SCA3 RAN translation of the expanded CAG repeats is initiated at upstream non-cognate codons. Schematic representation of *ATXN3* ORF-derived constructs: **a** AUG-78/32-117CAG and **c** 78/32-115CAG is shown with the indicated expanded CAG tracts (red, bold) and 5’ and 3’ flanking sequences (black, bold). HA-tag sequence is also indicated (green, bold). The identified canonical translation-dependent AUG start codon **(a)** and non-cognate RAN translation-dependent codons **(c)** are highlighted (light blue, bold). Amino acid sequences of AUG-initiated and RAN polyQ proteins are indicated in navy blue. **b** Representative LC-MS/MS spectra of the N-terminal part of the HA-immunoprecipitated polyQ proteins (^AC^SGTNLTSEELRK) from cells transfected with AUG-78/32-117CAG construct. **d, e** Representative LC-MS/MS spectra of the N-terminal part of the HA-immunoprecipitated RAN polyQ proteins (^AC^TSEELRK; ^AC^SEELRK) from cells transfected with 78/32-115CAG construct, indicating the RAN translation initiation sites

Interestingly, our proteomic analysis revealed that the canonically translated polyQ proteins from AUG-78/32-117CAG control construct had an initial acetylated serine, as revealed by both LysC and trypsin digestion, i.e., the obtained peptide sequence was ^AC^SGTNLTSEELRK (Fig. 5a and b, Fig. S8a). No peptides from polyQ proteins initiated from the canonical AUG codon contained N-terminal methionine, indicating that, although the first AUG codon is indeed decoded by an initiator Met-tRNA (iMet-tRNA), the incorporated methionine is removed rapidly by methionine aminopeptidases (MetAPs) followed by the efficient acetylation of the novel protein N-terminus by an N-terminal acetyltransferase (NAT) complex (Fig. 5b, Fig. S8a). Our observation is consistent with finding from previous studies showing that the complete cleavage of iMet by MetAPs is achievable for the following substrates: Met-Ser-, Met-Thr-, Met-Gly-, Met-Ala-, Met-Val-, Met-Cys- and Met-Pro- [55, 56]. Furthermore, the resulting new N-terminus (excluding Pro) was shown to be very efficiently Nt-acetylated by a major NAT complex – the NatA complex [57–59]. Since the Nt-acetylome is estimated to include as many as 80–90% of soluble human proteins, it provides a reliable indication of where translation initiation starts [59, 60].

In the case of the RAN polyQ proteins expressed in the HEK293T cells transfected with the 78/32-115CAG construct, we identified unique N-terminally acetylated peptides, ^AC^TSEELRK (2 peptides) and ^AC^SEELRK (2 peptides), that indicated the sites of RAN translation initiation (Fig. 5c–e, Fig. S9a). These data, obtained for both LysC and trypsin digestion, suggest that, similar to the proteins synthesized by canonical translation, the initial amino acid incorporated into this RAN polyQ protein is methionine, which is removed by MetAPs followed by N-terminal acetylation of the second amino acid (Thr or Ser) by NatA complex. Interestingly, our results strongly suggest that the expanded CAG repeats in *ATXN3* can induce RAN translation initiation at neighboring non-cognate codons, i.e., CUU and ACU (Fig. 5c-e, Fig. S9a). Moreover, the heterogeneity of the RAN translation initiation events may not be limited to only these two favorable codons, as some indications of RAN translation initiation upstream and downstream of the CUU and ACU codons have also been indicated by LC-MS/MS analysis (Fig. S9a) and previous RAN translation analysis based on constructs with shorter 5’ flanking sequences, e.g., 45/32-110CAG and 33/32-119CAG constructs (Fig. 4, Fig. S5).

In the case of RAN polyQ proteins isolated from the HEK293T cells transfected with the 33/32-119CAG construct, we failed to identify peptides showing RAN translation initiation site(s) upstream of the expanded CAG repeats, most likely because of the suboptimal peptide size (<5 amino acids) generated after LysC and trypsin digestion, which makes them difficult to identify in complex mixtures and through database searching (Fig. S10a). Similarly, in the case of all the RAN polyA proteins analyzed by LC-MS/MS, we were not able to identify, with high confidence, the location of RAN translation initiation (Fig. S8b, Fig. S9b, Fig. S10b). Based on our previously described experiments showing the extensive heterogeneity of RAN polyA protein sizes and because of the naturally occurring stop codon only eight amino acids upstream of the expanded repeats in *ATXN3*, we propose that the vast majority of the RAN polyA proteins were initiated within repeats at the GCA codons.

### Cellular stress enhances SCA3 RAN translation of the CAG repeats

In response to diverse stress stimuli such as hypoxia, viral infections or the presence of misfolded proteins, eukaryotic cells activate a cytoprotective signaling pathway termed the integrated stress response (ISR) to restore cellular homeostasis. The core event of ISR is phosphorylation of the α subunit of eIF2, which leads to global protein synthesis attenuation accompanied by the induction of alternative mechanisms of translation initiation of selected mRNAs [61–64]. Recently, it has been reported that RAN translation at CGG and GGGGCC repeats is enhanced by ISR activation [26, 27, 29, 36]. To examine whether this phenomenon can also occur in SCA3, cells transfected with selected RAN constructs (78/32-115CAG, 45/32-110CAG and 33/32-119CAG) were additionally exposed for 8 h to thapsigargin (TG), which causes endoplasmic reticulum (ER) stress, or to sodium arsenite (SA), which induces oxidative stress. As expected, both stressors led to increased eIF2α phosphorylation and upregulation of transcription factor C/EBP homologous protein (CHOP), one of the effectors of ISR, in both untransfected and transfected cells (Fig. 6a and b; Fig. S11a-d). Next, we found that treatment with either of these stress inducers significantly increased RAN translation for the polyA proteins in the cells expressing all the studied constructs (Fig. 6d, e, g and h, Fig. S11f and h). We also observed enhanced RAN translation for the polyQ proteins in cells treated with TG (Fig. 6c and e, Fig. S11e). In the case of oxidative stress, elevated levels of RAN polyQ proteins were observed only in the cells expressing the 78/32-115CAG and 45/32-110CAG constructs (Fig. 6f and h, Fig. S11g). In contrast to the RAN polyA proteins, treatment with SA did not increase the RAN polyQ levels in the cells expressing the 33/32-119CAG construct (Figure 6f and h).

**Fig. 6.**
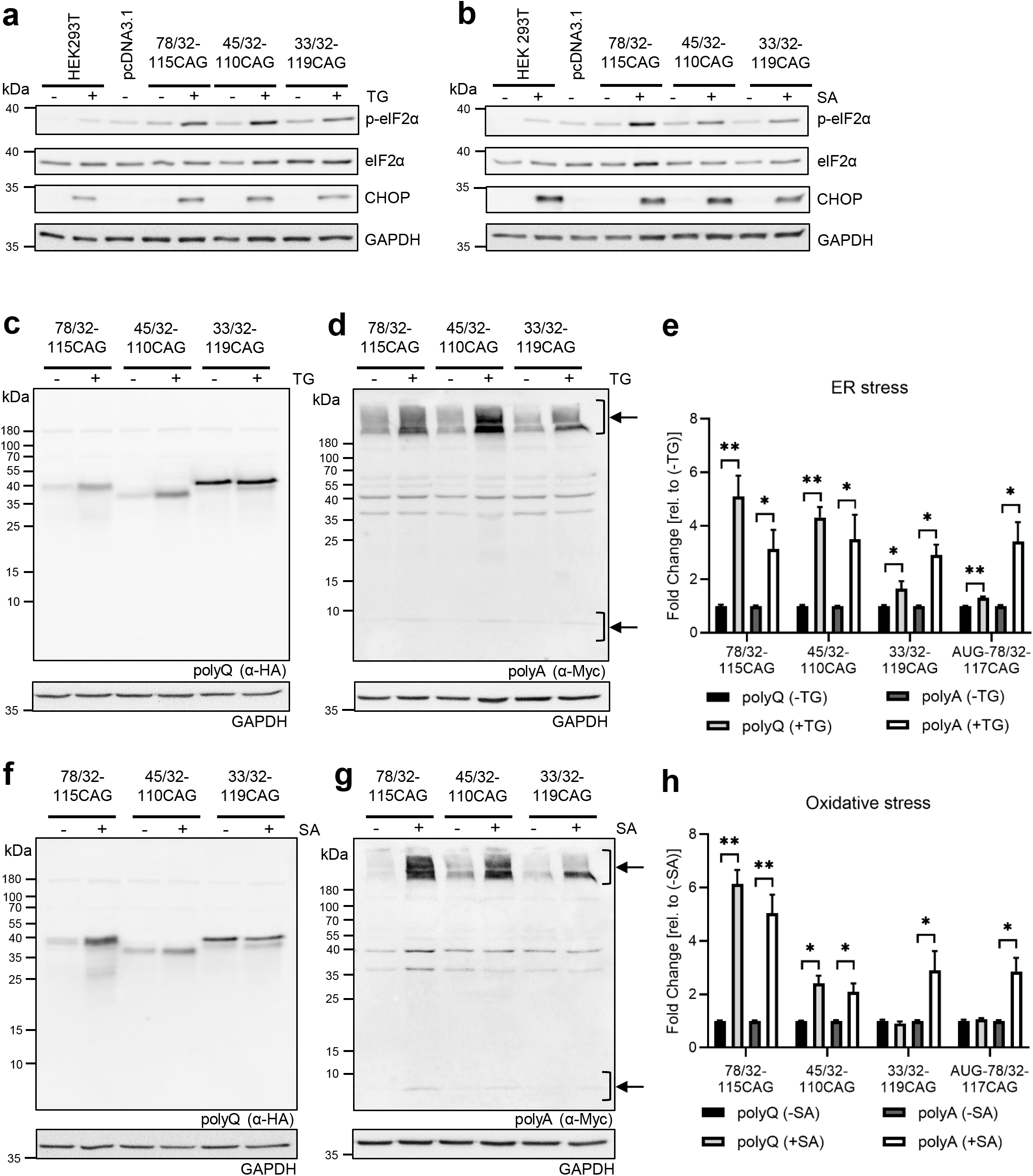
SCA3 RAN translation of the CAG repeats is upregulated upon cellular stress. For all the analyses presented in this figure, lysates were obtained from HEK293T cells 20 h after their transfection with the indicated constructs followed by 8 h of treatment with 1 μM thapsigargin (TG) or 50 μM sodium arsenite (SA). **a, b** Western blot analysis of phospho-eIF2α at the Ser51 site, eIF2α and CHOP in the HEK293T cells that were either transfected or not with the indicated constructs and treated or not with TG or SA. **c, d** Expression of the RAN polyQ and RAN polyA proteins after treatment or not with TG was analyzed by western blotting using anti-HA and anti-Myc antibodies, respectively. **e** Relative expression of the RAN polyQ and RAN polyA proteins and AUG-initiated polyQ proteins upon ER stress induction by TG and normalized to that of the untreated cells. In Fig. S11e and f, western blot analysis of the AUG-initiated polyQ, RAN polyQ and RAN polyA proteins in the HEK293T cells transfected with the AUG-117CAG and 45/32-110CAG constructs, and treated or not with TG is shown. The graph bars represent the mean value ± SEM from 5 biological replicates. Two-tailed *t* test, *p<0.05, **p<0.0005. **f, g** Expression of the RAN polyQ and RAN polyA proteins after treatment or not with SA was analyzed by western blotting using anti-HA and anti-Myc antibodies, respectively. **h** Relative expression of the RAN polyQ and RAN polyA proteins and AUG-initiated polyQ proteins upon oxidative stress induction by SA and normalized to that of the untreated cells. In Fig. S11g and h, western blot analysis of the AUG-initiated polyQ, RAN polyQ and RAN polyA proteins in the HEK293T cells transfected with the AUG-117CAG and 45/32-110CAG constructs, and treated or not with SA. The graph bars represent the mean value ± SEM from 5 biological replicates. Two-tailed *t* test, *p<0.05, **p<0.001. Specific RAN polyA products used in the quantitative analysis are indicated by black arrows. For all the western blot analyses, GAPDH was used as a loading control

As many cellular stressors negatively interfere with cap-dependent translation, we also studied the effect of ER and oxidative stress on the translation in cells transfected with AUG-78/32-117CAG construct. It is worth noting that, in addition to AUG-initiated polyQ proteins and RAN polyA proteins, RAN polyQ proteins could also be produced in these cells. In the cells transfected with the AUG-78/32-117CAG plasmid, the levels of eIF2a phosphorylation and CHOP were also markedly elevated under stress conditions (Fig. S11a-d). The administration of TG to cells expressing the AUG-78/32-117CAG construct caused a slight increase in polyQ protein levels (Fig. 6e, Fig. S11e), in contrast to the SA treatment, for which no such change was observed (Fig. 6h, Fig. S11g). Importantly, both treatments substantially elevated RAN translation for polyA proteins in the cells transfected with the AUG-78/32-117CAG plasmid (Fig. 6e and h; Fig. S11f and h). Thus, these data show that ISR can also enhance the production of RAN proteins at CAG repeats in SCA3.

## DISCUSSION

The discovery of RAN translation has revolutionized not only the understanding of the mechanisms underlying repeat expansion disorders but has also indicated novel therapeutic targets to combat these devastating diseases. To date, this unconventional translation has been reported in eight distinct diseases, including SCA8, DM1, c9ALS/FTD, FXTAS, HD, SCA31, DM2 and FECD, and a growing body of evidence clearly indicates that RAN translation products cause cellular toxicity [37–41]. Generally, RAN translation has been found to occur mainly across large repeat expansions in non-coding regions of the transcripts. Many reports have shown that these expanded repeats in RNA form a very stable secondary structure that is required for RAN translation [65–67]. It has been reported that mutant CAG repeats in polyQ diseases can also fold into stable hairpins, which might suggest that these repeats can also trigger RAN translation [66]. However, because of the location of these CAG repeats within open reading frames, studying RAN translation in polyQ diseases is technically challenging. To date, only Banez-Coronel and coworkers have provided strong evidence that this unusual translation can occur at CAG repeats in coding regions [15]. They showed the accumulation of four novel RAN proteins in HD human brain samples, which may be implicated in disease pathology.

Here, we report the first comprehensive study showing that RAN translation occurs at expanded CAG repeats in the *ATXN3* context and can contribute to the pathogenesis of SCA3. Furthermore, we provide, for the first time, some insights into the RAN translation mechanism in SCA3, showing that, in addition to the pathological length of expansion, the sequences surrounding the repeat region play important roles in SCA3 RAN translation initiation and efficiency. Moreover, SCA3 RAN translation is enhanced by the activation of ISR pathways.

We demonstrated the existence of two SCA3 RAN-translated polyQ and polyA proteins in cells transfected with constructs containing elongated CAG repeat regions and *ATXN3* sequence context without ATG codons. During the study of RAN translation in SCA8 and DM1, this unusual translation mode was also suggested for SCA3; however, a very short (20 bp) 5’ flanking sequence of *ATXN3* was used [10]. In our study, we used longer sequences surrounding the expansion region (78 bp in the 5’ direction of the repeats and 32 bp in the 3’ direction of the repeats), as it has been demonstrated that both the 5’ and 3’ flanking sequences of *ATXN3* influence the CAG repeat structure, which is known to be important for RAN translation initiation. We previously showed that 3’ terminal CAG repeats pair with an *ATXN3*-specific region named a pseudo-repeat sequence located in the 5’ direction of the repeat tracts [68]. Another rationale for the length of flanking sequences was the occurrence of the last ATG codons upstream of the repeats and the presence of the first stop codon in the alanine reading frame downstream of the repeats. It is very possible that these selected *ATXN3* flanking regions represent the minimal SCA3 RAN-translational unit.

SCA3 RAN polyQ proteins, both in diffused form or as aggregates, exhibited cytoplasmic and nuclear localization, whereas diffused SCA3 RAN polyA proteins were found exclusively in the cytoplasm, which is consistent with previous data regarding the subcellular distribution of other proteins containing expanded glutamine or alanine tracts [10, 15, 69, 70]. It is worth mentioning that the expanded polyA tracts can serve as nuclear export signals, and the cytoplasmic accumulation of polyA-containing proteins is dependent on the length of the alanine tract [71]. Despite distinct cellular localization, SCA3 RAN polyQ and polyA proteins are frequently found in the same cell, which suggests not only the common mechanism for RAN translation in various frames in SCA3 but also the cumulative toxic effect of these two distinct proteins in cells. We were not able to detect RAN polyS proteins, which may be explained by not having the optimal sequence and/or structure context to enable efficient RAN translation from this reading frame.

To date, an enormous effort has been made to better recognize the role of RAN proteins in the pathogenesis of repeat expansion disorders. Using various cellular and animal models it has been shown that RAN translation affects proteasome activity and nucleocytoplasmic transport, activates oxidative and ER stress, impairs protein translation and stress granule dynamics and/or induces cell death [37–41]. Here, we demonstrate that SCA3 RAN polyQ and RAN polyA proteins, similar to the RAN proteins described for other repeat expansion diseases, can cause cellular toxicity. First, these RAN polyQ proteins form aggregates, which are common pathological features of most neurodegenerative diseases and are considered potentially harmful to cells [72–74]. Second, in the cells expressing SCA3 RAN proteins, apoptosis is induced. Finally, the presence of SCA3 RAN proteins is associated with changes in nuclear envelope morphology, as determined by nuclear pore complex staining. The last finding is particularly important because it shows, for the first time, that RAN translation of CAG repeats might affect nuclear membrane integrity, potentially leading to dysfunction of nucleocytoplasmic transport. It was previously shown using i.a. HD mouse models that mutant huntingtin disrupts the nuclear envelope architecture and nucleocytoplasmic transport [50, 51]. There are also several reports indicating that nucleocytoplasmic transport is also impaired by RAN proteins [51, 75–79]. Moreover, interaction of RAN polyglycine (polyG) proteins translated from CGG repeats in FXTAS with the nuclear lamina protein LAP2β has been reported, which is associated with disorganization of the nuclear lamina structure [25]. Alterations in the nuclear lamina are also induced by dipeptide repeat proteins produced from GGGGCC repeats [75, 80]. The precise role of SCA3 RAN proteins in disease pathogenesis requires further investigation; however, it is very likely that the toxicity of these proteins could be associated with nuclear dysfunction, such as an impaired nuclear envelope and aberrant nucleocytoplasmic transport, leading to cell death.

Consistent with previous work on other repeat expansions, RAN translation of CAG repeats in the *ATXN3* sequence context is also strongly influenced by repeat length [20–23]. We did not observe RAN products translated at a nonpathogenic number of CAG repeats, and the amount of both RAN polyQ and RAN polyA proteins substantially increased with longer CAG expansion tracts. Moreover, we revealed an important role of the sequences surrounding the repeat region on SCA3 RAN translation initiation and efficiency. We showed that truncating the 5’ flanking sequence to 24 nt prevented RAN translation only in the glutamine frame, which suggested translation initiation upstream of the repeats, whereas the length of the downstream sequence affected RAN polyA proteins expression. Using mass spectrometry analysis, we confirmed that the synthesis of RAN polyQ proteins, in fact, is initiated at sequence preceding the CAG repeats. In the case of the RAN polyA proteins, we were not able to detect peptides indicating the initiation upstream of the repeats, which supports the hypothesis that translation in the alanine frame occurs within the repeats [10, 11]. Previous research on RAN translation in SCA8, FXTAS or c9ALS/FTD has provided evidence that the length and the content of the 5’ flanking sequences influence the initiation of RAN translation in specific reading frames, whereas sequences located downstream of the repeats can affect RAN translation in SCA2 [10, 11, 24–26, 28–30, 81]. Insertion of a stop codon in the *ATXN8* sequence immediately before the CAG repeats impedes RAN translation in the glutamine frame but not in the alanine or serine frames [10]. Similarly, shortening the *FMR1* 5’ UTR sequence located upstream of the CGG repeats or introducing stop codons at specific sites in this sequence inhibits the expression of RAN polyG but not RAN polyA proteins [11, 24, 25]. Further studies have shown that RAN polyG proteins are required for the initiation at an ACG near-cognate codon located 32 nt upstream of the CGG repeats [25]. Additionally, it has been revealed that RAN translation of GGGGCC repeats occurs at the near-cognate CUG codon located 24 nt before the repeats, and deletion of the sequence containing this codon abolishes translation [26, 28–30, 81]. The *ATXN3* 5’ flanking sequence used in our research contains two near-cognate AAG codons in the glutamine frame, one located 45 nt and the second 9 nt in the 5’ direction of the repeats. The presence of these codons could explain the appearance of two bands of RAN polyQ proteins, as well as the lack of expression of these proteins when the upstream sequence is shortened. However, the mutation of one of these codons (45 nt upstream) and analysis of the RAN translation of the 33/32 construct have shown that this codon is not involved in the translation initiation for the RAN polyQ proteins. Moreover, our mass spectrometry analysis revealed that the SCA3 RAN translation in the glutamine frame is initiated at the neighboring non-cognate CUU and ACU codons. The observed results could be explained by the fact that the AAG codon is one of the least efficient near-cognate start codons; moreover, in the *ATXN3* sequence, it is in a poor Kozak context [53, 54]. This finding is in agreement with proteomic studies showing that ACG and CUG codons are indeed initiation sites in the RAN translation of CGG and GGGGCC repeats, respectively [25, 30]. Generally, these codons appear to be the most efficient near-cognate start codons, and they are embedded in a good Kozak consensus sequence in causative genes for FXTAS and c9ALS/FTD [25, 30, 53, 54]. Based on previous studies concerning structural requirements for translation initiation, we initially assumed, that a stable hairpin formed by expanded CAG repeats downstream from the AAG codons would increase the likelihood of initiation at these sites [82–85]. Why does RAN translation in the glutamine frame initiate from CUU and ACU codons in a weak Kozak motif? It is likely that the structural context of these non-cognate codons, together with the downstream hairpin structure formed by the CAG repeats, makes these sites more accessible to the ribosome for the initiation of translation compared to the sites of the near-cognate AAG codons. It has been previously shown that structural RNA elements in which the initiation AUG codon is embedded have a strong impact on translation i.e., placing an AUG codon within a highly stable hairpin motif substantially reduced translational efficiency [86–90]. Nevertheless, despite the initiation at CUU and ACU codons that encode leucine and threonine, respectively, our data indicate that methionine is likely to be the initial amino acid incorporated into the RAN polyQ proteins. However, due to the specific amino acids in the second position, this methionine is completely removed by MetAPs, and the next amino acid, threonine or serine, is acetylated by NAT complex [55–59]. Previous mass spectrometry analysis of proteins produced through RAN translation of CGG and GGGGCC repeats also revealed that, despite imperfect match methionine was the initial amino acid for polyG translated from the ACG codon and for glycine-alanine dipeptide repeat proteins translated from the CUG codon [25, 30]. In contrast to the results of our analysis, due to the presence of glutamic acid in the second position, methionine could not be excised and therefore was detected in the identified peptides [25, 30]. N-terminal methionine was also not observed in the peptides derived from SCA8 RAN polyA proteins, which was explained as a potential consequence of the removal by a specific peptidase or an initiation mechanism without a requirement for methionine [10]. In conclusion, our findings suggest that RAN translation can also be initiated from non-cognate codons incorporating methionine as the first amino acid.

In the present study, the occurrence of RAN translation across a full-length *ATXN3* transcript was also examined. We used constructs expressing, in addition to different lengths of CAG repeats, two variants of *ATXN3* cDNA: one canonically translated because of the presence of an ATG start codon and the second one in which stop codons were introduced to abolish conventional ATXN3 translation. Regardless of the constructs used, including those containing expanded CAG repeats, we were not able to observe RAN proteins. This could be due to the fact that RAN translation of full-length mutant *ATXN3* mRNA occurs but at undetectable level until the toxicity-inducing events favoring this unconventional mode of translation successively accumulate as a result of disease progression. We cannot also exclude the possibility that SCA3 RAN translation might occur across at not yet identified, shorter and more pathogenic *ATXN3* isoforms, such as the transcripts used in our study, resulting from aberrant splicing. To date, 56 alternative splicing variants of *ATXN3* have been described [91]. In our study, we used the transcript containing 11 exons that encode the predominant *ATXN3* isoform in human and murine brain tissue and an additional SNP (CGG instead GGG) immediately downstream of the CAG repeat, which is known to be associated with SCA3 [92–95]. As the presence of a CAG expansion sequence in the SCA3 YAC mouse model is associated with increased level of alternatively spliced *ATXN3* isoform lacking exon 11, it is possible that such transcript could be more susceptible to RAN translation [96]. It is worth noting that, dependent on expanded CAG repeats, deregulated splicing of *HTT* exon 1 has been shown to lead to the production of short polyadenylated mRNA encoding a highly toxic HTT fragment [97, 98]. However, it remains unexplained which transcript in HD patients is indeed RAN translated: the normally processed full-length *HTT*, the aberrantly spliced small mRNA or both. Further research on RAN translation in SCA3 is required, including an analysis of the presence of RAN proteins in patient brain tissues.

It is commonly known that, in response to stress conditions, such as ER stress from misfolded proteins, oxidative stress or viral infections, the ISR pathway is activated, leading to the phosphorylation of eIF2α and causing global cap-dependent translation attenuation but upregulation of the translation of certain mRNAs in an unconventional manner [61–64]. Recent mechanistic studies on RAN translation clearly indicate that cellular stress is another trigger for this non-canonical mode of protein synthesis. It has been reported that RAN translation of GGGGCC and CGG repeats is enhanced by diverse stress stimuli [26, 27, 29, 36]. Our results are consistent with these data and indicate that RAN translation of *ATXN3*-associated repeat expansion sequences can also be driven by ISR. We demonstrated enhanced production of both SCA3 RAN polyQ and RAN polyA proteins upon ER stress and oxidative stress, which was accompanied by increased levels of p-eIF2α and CHOP. In contrast to previous studies, we did not report the inhibition of canonical translation under stress conditions. The differences among studies might be related to the control of the AUG-initiated translation. In the previous studies, vectors containing nanoluciferase or firefly luciferase sequences with AUG codons were used to study canonical translation, whereas in our research, we used the construct AUG-78/32-117CAG, which, in addition to AUG codon and *ATXN3*-specific sequences, also contained the expanded CAG repeat region [26, 27, 29]. As we demonstrated that RAN polyA proteins can be produced from this construct and that the amount of these proteins significantly increased after cellular stress induction, it is very likely that, under these conditions, RAN translation in the glutamine frame was also upregulated. Thus, we assume that the amount of polyQ proteins observed for the cells expressing the AUG-78/32-117CAG construct under ER and oxidative stress was the result of reduced canonical translation and increased RAN translation. Altogether, as age-dependent accumulation of misfolded mutant ataxin-3 can lead to activation of various cellular stress pathways, it is very likely that the amount of RAN proteins increase over time and, as a consequence, the pathogenesis of SCA3 is accelerated [1–4].

Overall, we provide new insights into the SCA3 RAN translation mechanism and pathogenesis, which may constitute possible novel targets of great therapeutic value in SCA3. In addition, our findings broaden our understanding of the requirements for RAN translation in general, particularly indicating the important role of the sequences surrounding an expansion region and the possibility of initiation from non-cognate codons.

## ACKNOWLEDGEMENTS

This work is dedicated to the memory of Professor Wlodzimierz J. Krzyzosiak. We thank Professor Marta Olejniczak from Department of Genome Engineering for reviewing the manuscript and every day stimulating discussions. We thank Professor Jerzy Ciesiolka from Department of RNA Biochemistry for helpful suggestions. The confocal microscopy analyses were performed in the Laboratory of Subcellular Structures Analysis, Institute of Bioorganic Chemistry, Polish Academy of Sciences, Poznan. Proteomic mass spectrometry analyses were performed in the Mass Spectrometry Laboratory, Institute of Biochemistry and Biophysics, Polish Academy of Sciences, Warsaw. This work was supported by the National Science Centre, Poland [2015/19/D/NZ5/02183 to M.J.-C., 2012/06/A/NZ1/00094 to W.J.K., and 2015/19/B/NZ2/02453 to A.C.]; the Polish Ministry of Science and Higher Education (statutory funds for young investigators to M.J.-C.); and the Polish Ministry of Science and Higher Education under the KNOW program [01/KNOW2/2014].

## ABBREVIATIONS

ATXN3: ataxin-3
c9ALS/FTD: c9orf72 amyotrophic lateral sclerosis/frontotemporal dementia
CHOP: C/EBP homologous protein
DM1: myotonic dystrophy type 1 and type 2
ER: endoplasmic reticulum
FECD: Fuchs’ endothelial corneal dystrophy
FXTAS: fragile X tremor ataxia syndrome
iMet-tRNA: initiator Met-tRNA
ISR: integrated stress response
MetAPs: methionine aminopeptidases
NAT: N-terminal acetyltransferase
NPC: nuclear pore complex
polyA: polyalanine
polyQ: polyglutamine
polyS: polyserine
RAN translation: repeat associated non-ATG translation
SA: sodium arsenite
SCA3: spinocerebellar ataxia type 3
TG: thapsigargin

## DECLARATIONS

### Ethics approval

Not applicable.

### Consent to participate

Not applicable.

### Consent for publication

Not applicable.

### Availability of data and material

All data generated or analyzed during this study are included in this published article and its supplementary information files.

### Code availability

Not applicable.

### Authors’ contributions

Authors contribution: M.J.-C. designed, coordinated and performed the research. A.C. designed and performed immunoprecipitation analysis and analyzed mass spectrometry data. A.A.K. and M.O.U.-T. assisted in some of the experiments. W.J.K. and A.F. supervised the study. M.J.-C. wrote the original draft with exception of mass spectrometry results which were written by A.C.. M.J-C., A.C. and A.F. reviewed and edited the manuscript with editorial input from A.A.K. and M.O.U.-T. All authors read and approved the final manuscript.

### Corresponding authors

Correspondence to Agnieszka Fiszer and Magdalena Jazurek-Ciesiolka

