## Supplementary Figures for "RAN translation of the expanded CAG repeats in the SCA3 disease context"

### Fig. S1

**a**

(…)**ATG**GAGTCCATCTTCCACGAGAAACAAGAAGGCTCACTTTGTGCTCAACATTGCCTGAATAACTTATTGCA
AGGAGAATATTTTAGCCCTGTGGAATTATCCTCAATTGCACATCAGCTGGATGAGGAGGAGAGGATGAGAATG
GCAGAAGGAGGAGTTACTAGTGAAGATTATCGCACGTTTTTACAGCAGCCTTCTGGAAATATGGATGACAGTG
GTTTTTTCTCTATTCAGGTTATAAGCAATGCCTTGAAAGTTTGGGGTTTAGAACTAATCCTGTTCAACAGTCCA
GAGTATCAGAGGCTCAGGATCGATCCTATAAATGAAAGATCATTTATATGCAATTATAAGGAACACTGGTTTAC
AGTTAGAAAATTAGGAAAACAGTGGTTTAACTTGAATTCTCTCTTGACGGGTCCAGAATTAATATCAGATACAT
ATCTTGCACTTTTCTTGGCTCAATTACAACAGGAAGGTTATTCTATATTTGTTGTTAAGGGTGATCTGCCAGATT
GCGAAGCTGACCAACTCCTGCAGATGATTAGGGTCCAACAGATGCATCGACCAAAACTTATTGGAGAAGAATT
AGCACAACTAAAAGAGCAAAGAGTCCATAAAACAGACCTGGAACGAATGTTAGAAGCAAATGATGGCTCAGGA
ATGTTAGACGAAGATGAGGAGGATTTGCAGAGGGCTCTGGCACTAAGTCGCCAAGAAATTGACATGGAAGAT
GAGGAAGCAGATCTCCGCAGGGCTATTCAGCTAAGTATGCAAGGTAGTTCCAGAAACATATCTCAAGATATGA
CACAGACATCAGGTACAAATCTTACTTCAGAAGAGCTTCGGAAGAGACGAGAAGCCTACTTTGAAAAACAGCA
GCAAAAGCAGCAA(CAGCAGCAGCAG)nCGGGACCTATCAGGACAGAGTTCACATCCATGTGAAAGGCCAGCC
ACCAGTTCAGGAGCACTTGGGAGTGATCTAGGTGATGCTATGAGTGAAGAAGACATGCTTCAGGCAGCTGTG
ACCATGTCTTTAGAAACTGTCAGAAATGATTTGAAAACAGAAGGAAAAAAA**TAA**(…)

**b**

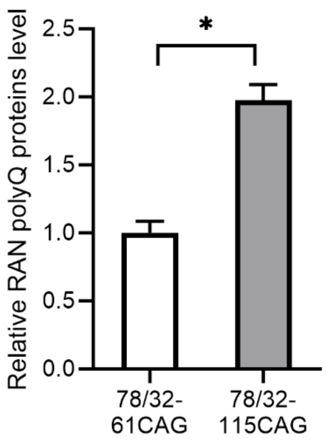


**c**

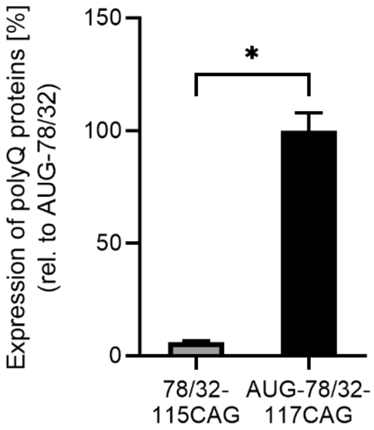


**Fig. S1 a** Many alternative ATG codons are present within the *ATXN3* sequence. The coding *ATXN3* sequence obtained from SCA3 patient-derived fibroblasts (GM06153) is given. The ATG start codon and TAA stop codon are in bold. The position of the alternative ATG codons in various frames, upstream of the repeated tract, is marked in blue, while the position of the stop codons for the alanine and serine frames, downstream of the repeat tract, is marked in green. The flanking sequence used in the series of 78/32 constructs is underlined. **b** Relative expression of RAN polyQ proteins from 78/32-61CAG and 78/32-115CAG constructs. **c** Relative expression of RAN polyQ proteins from the 78/32-115CAG construct normalized to the AUG-78/32-117CAG control. The graph bars represent the mean value ± SEM from 5 biological replicates. Two-tailed *t* test, *p<0.0002

Fig. S2

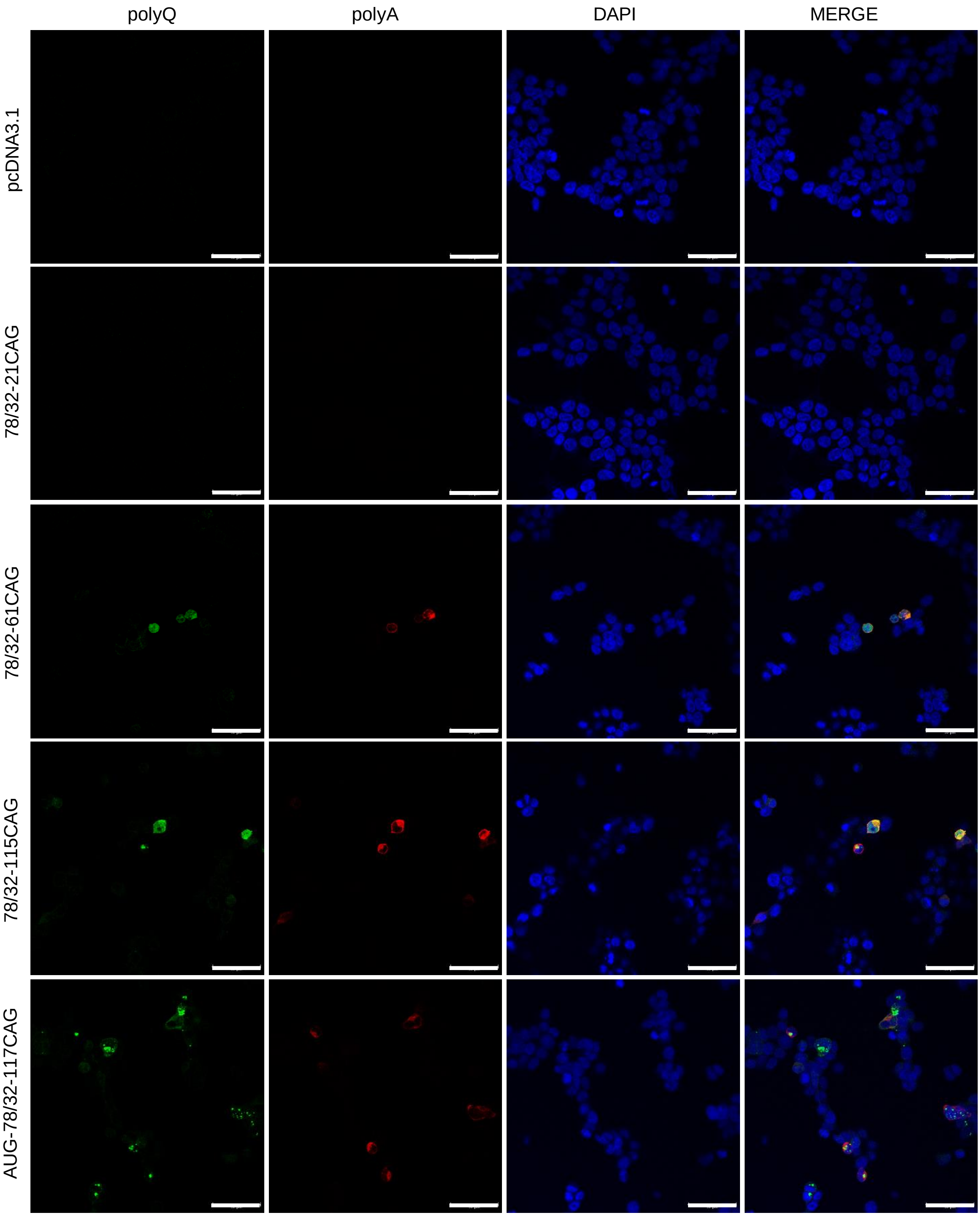
polyQ
polyA
DAPI
MERGE
pcDNA3.1
78/32-21CAG
78/32-61CAG
78/32-115CAG
AUG-78/32-117CAG

**Fig. S2** Cellular localization of SCA3 RAN polyQ and RAN polyA proteins. Representative images of immunofluorescence staining for the AUG-initiated and RAN polyQ proteins (green signal, anti-HA antibody) and RAN polyA proteins (red signal, anti-Myc antibody) in the HEK293T cells 48 h after transfection with the indicated constructs. DAPI (blue signal) was used to stain nuclei. Scale bar=50 μm

#### Fig. S3

**a**

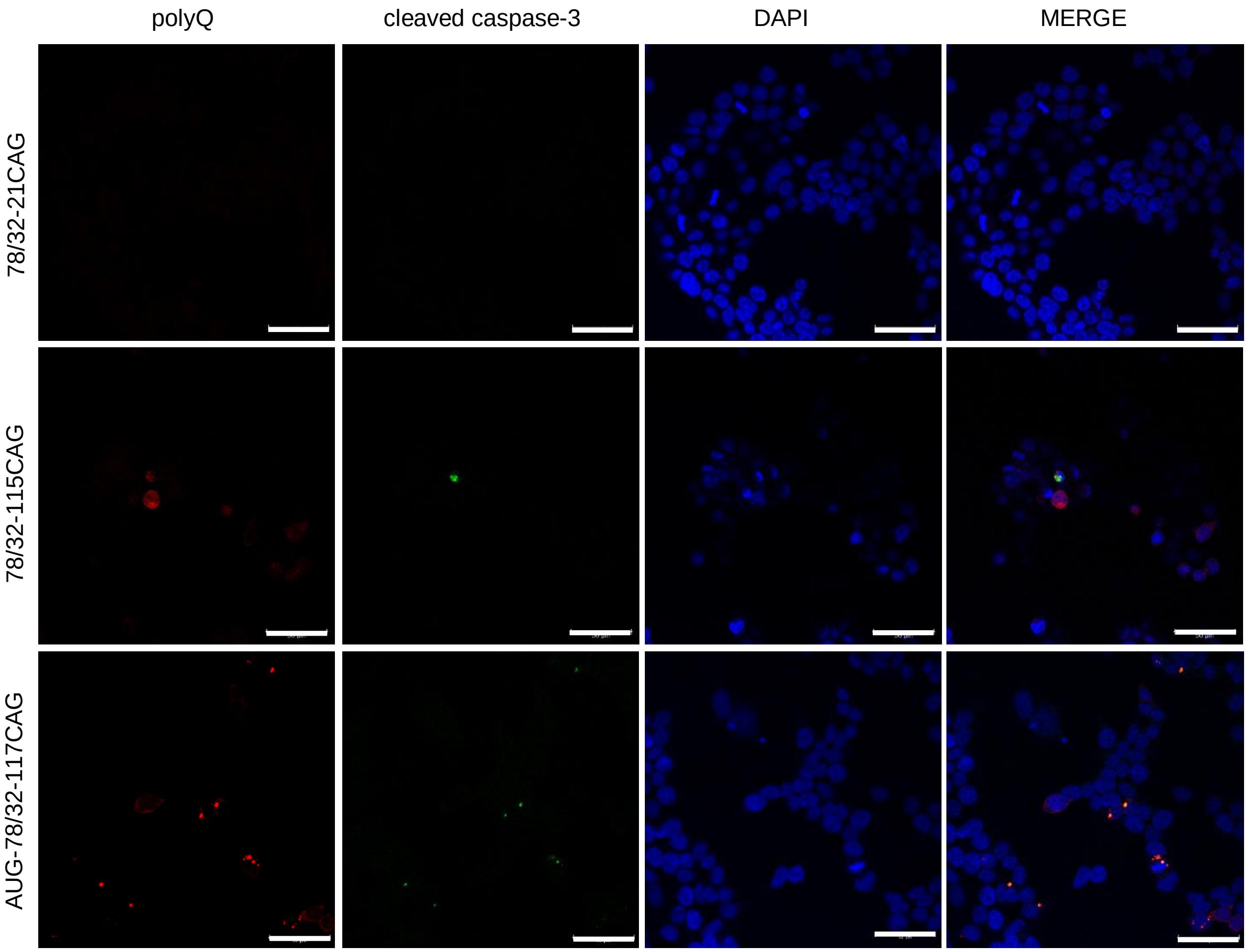
polyQ
cleaved caspase-3
DAPI
MERGE
78/32-21CAG
78/32-115CAG
AUG-78/32-117CAG

**b**

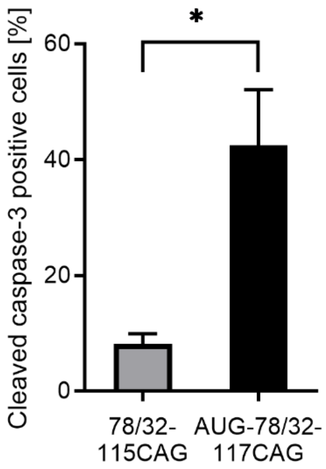
Cleaved caspase-3 positive cells [%]
60
40
20
0
*
78/32-115CAG
AUG-78/32-117CAG

**Fig. S3** SCA3 RAN translation induces cell death. **a** Representative images of immunofluorescence staining for AUG-initiated and RAN polyQ proteins (red signal, anti-HA antibody) and cleaved caspase-3 (green signal) in the HEK293T cells 48 h after transfection with the indicated constructs. DAPI (blue signal) was used to stain nuclei. Scale bar=50 μm. **b** Quantitative analysis of the cells expressing AUG-initiated or RAN polyQ proteins together in the presence of cleaved caspase-3; 100% represents cells expressing AUG-initiated or RAN polyQ proteins. The graph bars represent the mean value ± SEM from 3 biological replicates. Two-tailed *t* test, *p<0.01

#### Fig. S4

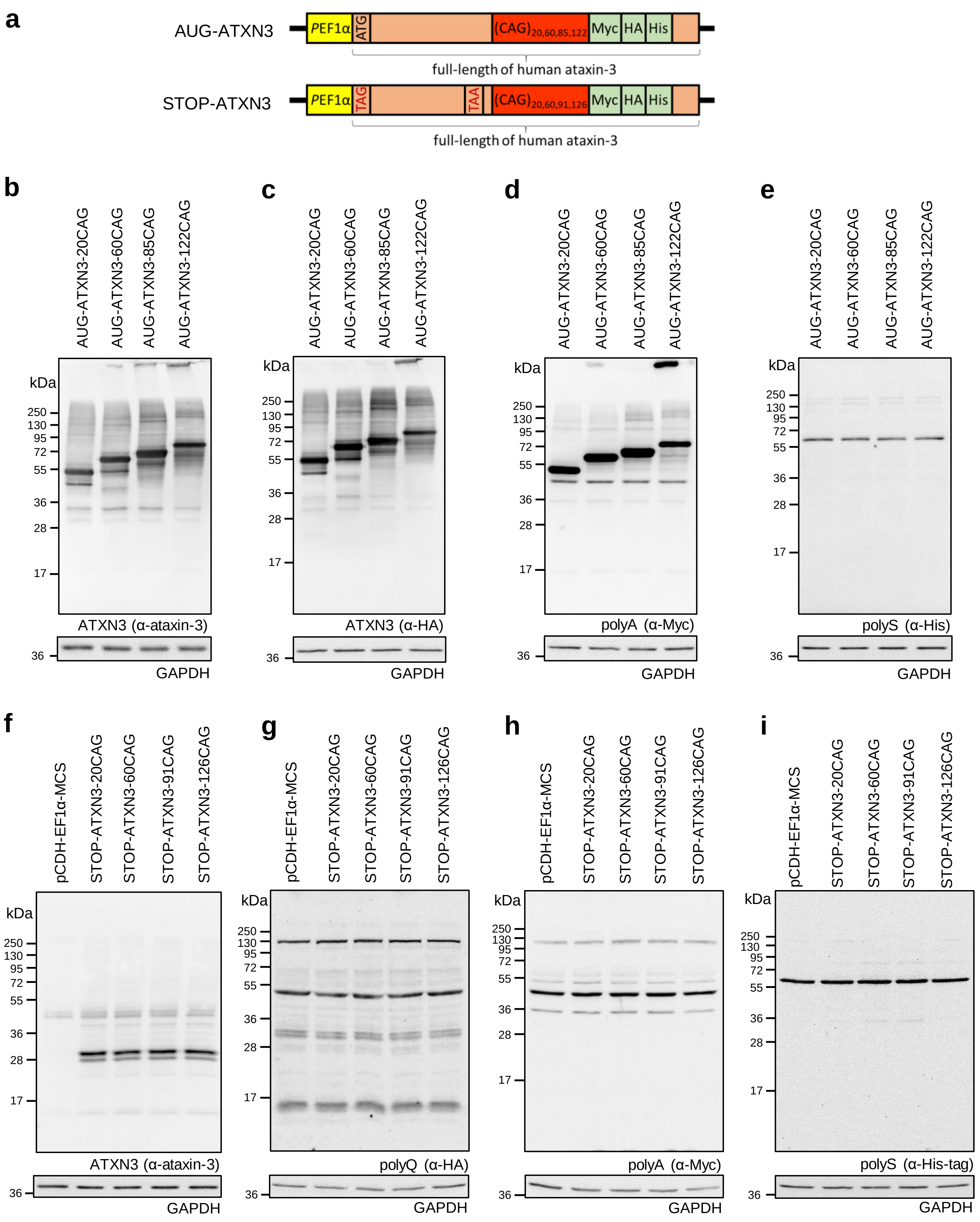

a
AUG-ATXN3
PEF1α
ATG
(CAG)20,60,85,122
Myc
HA
His
full-length of human ataxin-3
STOP-ATXN3
PEF1α
TAG
TAA
(CAG)20,60,91,126
Myc
HA
His
full-length of human ataxin-3
b
AUG-ATXN3-20CAG
AUG-ATXN3-60CAG
AUG-ATXN3-85CAG
AUG-ATXN3-122CAG
kDa
250
130
95
72
55
36
28
17
ATXN3 (α-ataxin-3)
36
GAPDH
c
AUG-ATXN3-20CAG
AUG-ATXN3-60CAG
AUG-ATXN3-85CAG
AUG-ATXN3-122CAG
kDa
250
130
95
72
55
36
28
17
ATXN3 (α-HA)
36
GAPDH
d
AUG-ATXN3-20CAG
AUG-ATXN3-60CAG
AUG-ATXN3-85CAG
AUG-ATXN3-122CAG
kDa
250
130
95
72
55
36
28
17
polyA (α-Myc)
36
GAPDH
e
AUG-ATXN3-20CAG
AUG-ATXN3-60CAG
AUG-ATXN3-85CAG
AUG-ATXN3-122CAG
kDa
250
130
95
72
55
36
28
17
polyS (α-His)
36
GAPDH
f
pCDH-EF1α-MCS
STOP-ATXN3-20CAG
STOP-ATXN3-60CAG
STOP-ATXN3-91CAG
STOP-ATXN3-126CAG
kDa
250
130
95
72
55
36
28
17
ATXN3 (α-ataxin-3)
36
GAPDH
g
pCDH-EF1α-MCS
STOP-ATXN3-20CAG
STOP-ATXN3-60CAG
STOP-ATXN3-91CAG
STOP-ATXN3-126CAG
kDa
250
130
95
72
55
36
28
17
polyQ (α-HA)
36
GAPDH
h
pCDH-EF1α-MCS
STOP-ATXN3-20CAG
STOP-ATXN3-60CAG
STOP-ATXN3-91CAG
STOP-ATXN3-126CAG
kDa
250
130
95
72
55
36
28
17
polyA (α-Myc)
36
GAPDH
i
pCDH-EF1α-MCS
STOP-ATXN3-20CAG
STOP-ATXN3-60CAG
STOP-ATXN3-91CAG
STOP-ATXN3-126CAG
kDa
250
130
95
72
55
36
28
17
polyS (α-His-tag)
36
GAPDH

**Fig. S4** RAN translation analysis of the cellular models expressing full-length *ATXN3*. **a** Diagram of the constructs containing canonically translated (upper panel, AUG-ATXN3 constructs) or non-canonically translated (lower panel, STOP-ATXN3 constructs) full-length cDNA of *ATXN3* with various lengths of CAG repeat sequences. In the STOP-ATXN3 constructs, an ATG codon was mutated to stop codon TAG and an additional stop codon TAA was introduced after the last ATG codon located upstream of the CAG repeat region. In all constructs just downstream of CAG repeats, Myc-, HA- and His-tags were placed in alanine, glutamine and serine reading frames, respectively. For all the analyses presented in this figure, lysates were obtained from HEK293T cells 48 h after their transfection with the indicated constructs. **b, c** Expression of the exogenous ATXN3 was analyzed by western blotting using anti-ATXN3 and anti-HA antibodies. **d , e** Expression of the RAN polyA and RAN polyS proteins was analyzed by western blotting using anti-Myc and anti-His antibodies, respectively. No specific signals for these proteins were detected. **f, g** Expression of the exogenous ATXN3 was analyzed by western blotting using anti-ATXN3 and anti-HA antibodies. Introducing stop codons abolished translation of exogenous ATXN3. **g, h, i** Expression of the RAN polyQ, RAN polyA and RAN polyS proteins was analyzed by western blotting using anti-HA, anti-Myc and anti-His antibodies, respectively. No specific signals for these proteins were detected. For all the western blot analyses, GAPDH was used as a loading control

### Fig. S5

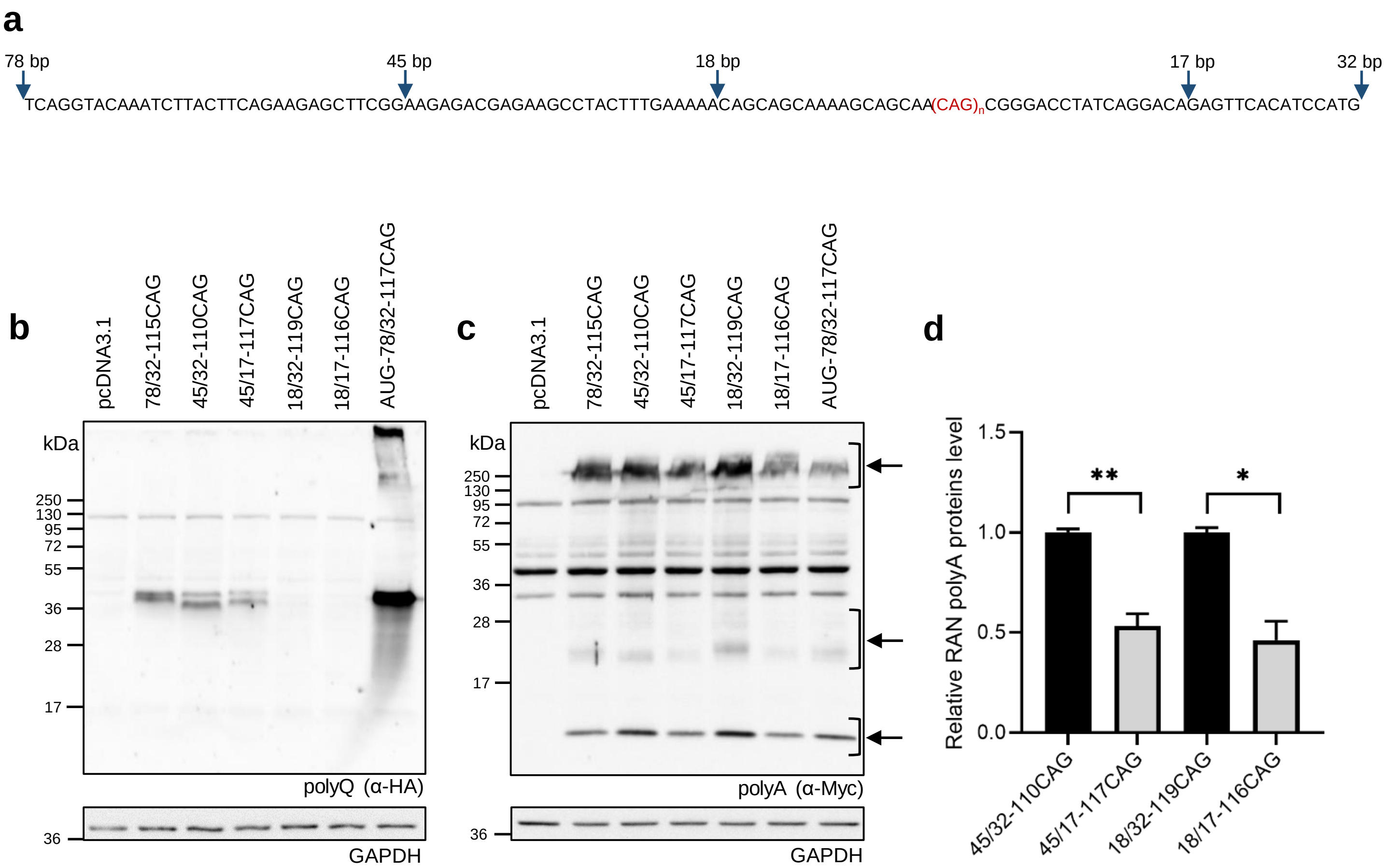


**Fig. S5** The effect of 5' and 3' *ATXN3* flanking sequences on the RAN translation efficiency of the CAG repeats. **a** *ATXN3* 5' and 3' flanking sequences of the 78/32-115CAG, 45/32-110CAG, 45/17-117CAG, 18/32-119CAG and 18/17-116CAG constructs. The blue arrows indicate the length of the flanking sequence upstream and downstream of the repeats used in particular RAN constructs. For all the analyses presented in this figure, lysates were obtained from HEK293T cells 48 h after their transfection with the indicated constructs. **b** Expression of the RAN polyQ proteins was analyzed by western blotting using an anti-HA antibody. **c** Expression of the RAN polyA proteins was analyzed by western blotting using an anti-Myc antibody. **d** Relative expression of the RAN polyA proteins from the indicated constructs normalized to that of the 45/32-110CAG or 18/32-119CAG constructs. Specific RAN polyA products used in the quantitative analysis are indicated by black arrows. The graph bars represent the mean value ± SEM from 4 biological replicates. Two-tailed t test, *p<0.002, **p<0.0005. For all the western blot analyses, GAPDH level was used as a loading control

**Fig. S6**

**a**

**78/32-115CAG**

UCAGGUACAAAUCUUACUUCAGAAGAGCUUCGGAAGAGACGAGAAGCCUACUUUGAAAAACAGCAGCAAAAGCAGCAA(CAG)n

**45/32-110CAG and 45/17-117CAG**

AAGAGACGAGAAGCCUACUUUGAAAAACAGCAGCAAAAGCAGCAA(CAG)$_n$

**18/32-119CAG and 18/17-116CAG**

CAGCAGCAAAAGCAGCAA(CAG)$_n$

AAG – polyQ (CAG); AGG/ACG – polyA (GCA); AUC/AAG/UUG – polyS (AGC)

AAG → CAG

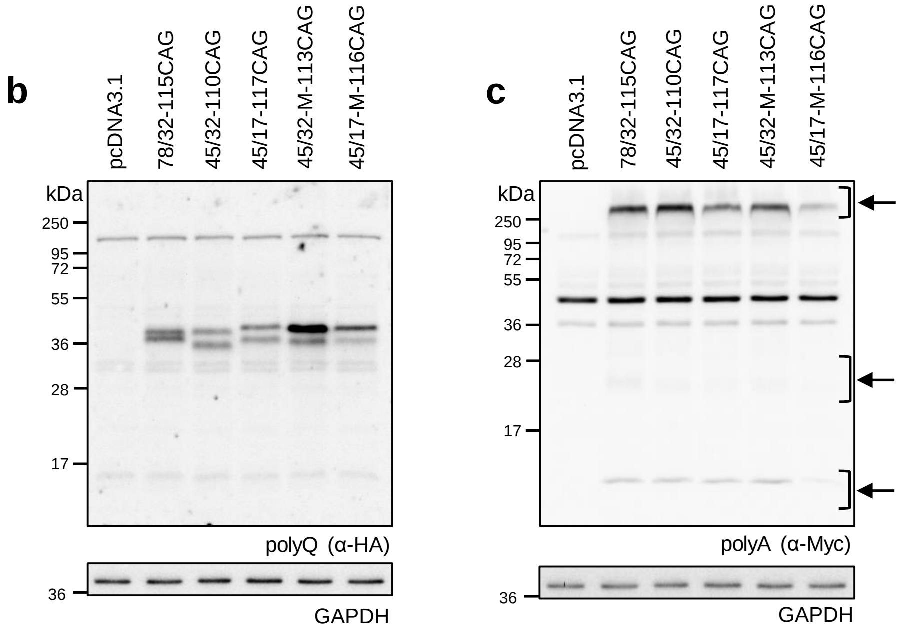


**Fig. S6** The AAG start codon is not involved in SCA3 RAN translation initiation in the glutamine frame. **a** The presence of AUG-like start codons in the 5' flanking sequence of *ATXN3* in the 78/32-115CAG, 45/32-110CAG, 45/17-117CAG, 18/32-119CAG and 18/17-116CAG constructs. The RNA sequence is given. Near-cognate codons in the glutamine frame are shown in violet; in the alanine frame are shown in blue; and in the serine frame are shown in green. The AAG codon mutated to CAG is underlined. For all the analyses presented in this figure, lysates were obtained from HEK293T cells 48 h after their transfection with the indicated constructs. **b** Expression of the RAN polyQ proteins was analyzed by western blotting using an anti-HA antibody. **c** Expression of the RAN polyA proteins was analyzed by western blot using an anti-Myc antibody. Specific products are indicated by black arrows. For all the western blot analyses, GAPDH was used as a loading control

**Fig. S7**

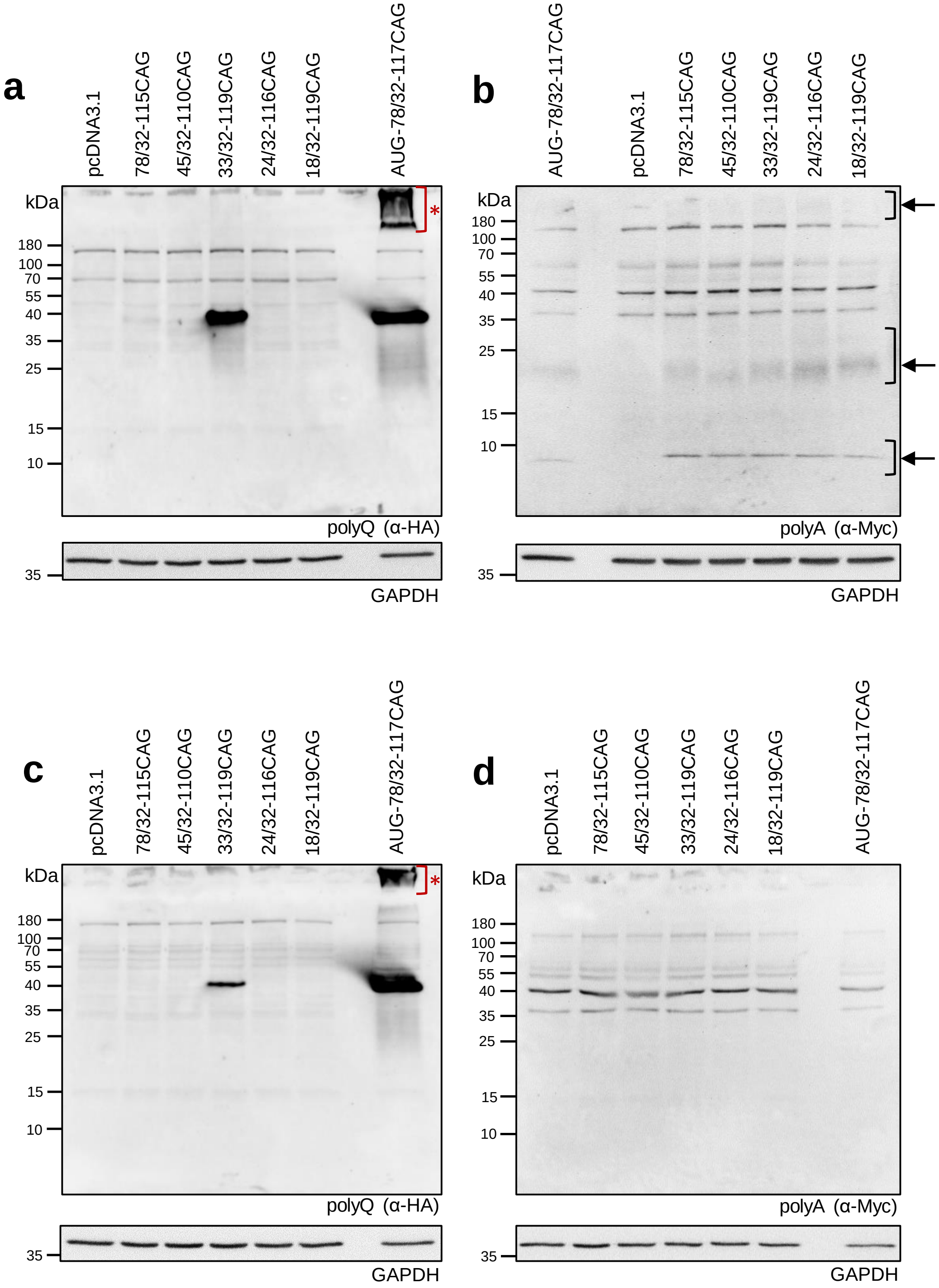
a
pcDNA3.1
78/32-115CAG
45/32-110CAG
33/32-119CAG
24/32-116CAG
18/32-119CAG
AUG-78/32-117CAG
kDa
180
100
70
55
40
35
25
15
10
*
polyQ (α-HA)
35
GAPDH
b
AUG-78/32-117CAG
pcDNA3.1
78/32-115CAG
45/32-110CAG
33/32-119CAG
24/32-116CAG
18/32-119CAG
kDa
180
100
70
55
40
35
25
15
10
polyA (α-Myc)
35
GAPDH
c
pcDNA3.1
78/32-115CAG
45/32-110CAG
33/32-119CAG
24/32-116CAG
18/32-119CAG
AUG-78/32-117CAG
kDa
180
100
70
55
40
35
25
15
10
*
polyQ (α-HA)
35
GAPDH
d
pcDNA3.1
78/32-115CAG
45/32-110CAG
33/32-119CAG
24/32-116CAG
18/32-119CAG
AUG-78/32-117CAG
kDa
180
100
70
55
40
35
25
15
10
polyA (α-Myc)
35
GAPDH

**Fig. S7** The efficiency of SCA3 RAN translation of the CAG repeats is dependent on the cell type. For all the analyses presented in this figure, lysates were obtained from HeLa or SH-SY5Y cells 48 h after their respective transfection with the indicated constructs. **a** Expression of the AUG-initiated and RAN polyQ proteins in HeLa cells was analyzed by western blotting using an anti-HA antibody. Aggregated AUG-initiated polyQ proteins are indicated by the red asterisk. **b** Expression of the RAN polyA proteins in the HeLa cells was analyzed by western blotting using an anti-Myc antibody. Specific products are indicated by black arrows. **c** Expression of the AUG-initiated and RAN polyQ proteins in the SH-SY5Y cells was analyzed by western blotting using an anti-HA antibody. Aggregated AUG-translated polyQ proteins are indicated by the red asterisk. **d** Expression of the RAN polyA proteins in the SH-SY5Y cells was analyzed by western blotting using an anti-Myc antibody. No specific products were detected. For all the western blot analyses, GAPDH was used as a loading control

**Fig. S8**

**a**

| Query | AUG-78/32-117CAG | | Ions Score | Expect |
|---|---|---|---|---|
| 799 | EAYFEK | - | 32 | 0,0006 |
| 3491 | REAYFEK | - | 45 | 0,0000 |
| 10246 | 'RREAYFEK | - | 33 | 0,0005 |
| 7874 | RREAYFEK | - | 31 | 0,0170 |
| 7872 | RREAYFEK | - | 31 | 0,0200 |
| 7875 | RREAYFEK | - | 40 | 0,0022 |
| 4469 | LTSEELRK | - | 38 | 0,0001 |
| 11614 | TNLTSEELRK | - | 39 | 0,0001 |
| 11613 | TNLTSEELRK | - | 34 | 0,0004 |
| 18834(D) | *SGTNLTSEELRK | - | 52 | 0,0000 |
| 18784 | *SGTNLTSEELRK | - | 50 | 0,0000 |
| 18783 | *SGTNLTSEELRK | - | 72 | 0,0000 |
| 18779 | *SGTNLTSEELRK | - | 74 | 0,0000 |
| 18780 | *SGTNLTSEELRK | - | 87 | 0,0000 |
| 13674 | GTNLTSEELRK | - | 41 | 0,0001 |
| 18871 | MGTNLTSEELRK | - | 38 | 0,0002 |
| | **MSGTNLTSEELRKRREAYFEKQQQKQQQQQQQQQQQQQQQQQQQQQQQ | | | |
| 15792(OD) | MGTNLTSEELR | - | 71 | 0,0000 |
| 20427 | *SGTNLTSEELRK | - | 57 | 0,0000 |
| 20428 | *SGTNLTSEELRK | - | 49 | 0,0000 |
| 20456(D) | *SGTNLTSEELRK | - | 58 | 0,0000 |
| 2328 | LTSEELR | - | 32 | 0,0007 |
| 6016 | 'REAYFEK | - | 30 | 0,0010 |
| 1076 | EAYFEK | - | 47 | 0,0005 |
| 4217 | REAYFEK | - | 40 | 0,0025 |
| 1076 | EAYFEK | - | 47 | 0,0005 |
| | QQQQQQQQQQQQQQQQQQQQQQQQQQQQQQQQQQQQQQQQQQQQQQQQQQQQQQQQQQQQ | | | |
| 26834(D) | DLSGQSSHPWNSNK | - | 40 | 0,0002 |
| 26815 | DLSGQSSHPWNSNK | - | 112 | 0,0000 |
| 33521(D) | RDLSGQSSHPWNSNK | - | 35 | 0,0003 |
| 33519(D) | RDLSGQSSHPWNSNK | - | 40 | 0,0001 |
| 33518(D) | RDLSGQSSHPWNSNK | - | 33 | 0,0006 |
| 33517(D) | RDLSGQSSHPWNSNK | - | 61 | 0,0000 |
| 33475 | RDLSGQSSHPWNSNK | - | 46 | 0,0000 |
| 33476 | RDLSGQSSHPWNSNK | - | 70 | 0,0000 |
| 35835 | 'RDLSGQSSHPWNSNK | - | 36 | 0,0002 |
| 42952(G) | QQRDLSGQSSHPWNSNK | - | 56 | 0,0000 |
| 62557(G) | QQQQQQRDLSGQSSHPWNSNK | - | 43 | 0,0001 |
| | QQQQQQQQQQQQQQQQQQQQQQQQQQQQQQQQQQQQRDLSGQSSHPWNSNKNSSQKRICY | | | |
| 59573(G) | QQQQQRDLSGQSSHPWNSNK | - | 72 | 0,0000 |
| 59649(GDDD) | QQQQQRDLSGQSSHPWNSNK | - | 34 | 0,0005 |
| 50694 | QQQQQQQQQQQQQQQR | - | 37 | 0,0002 |
| 80199(6xD) | LQQQQQQQQQQRDLSGQSSHPWNSNK | - | 32 | 0,0007 |
| 50903(D) | DLSGQSSHPWNSNKNSSQK | - | 71 | 0,0000 |
| 28530 | DLSGQSSHPWNSNK | - | 108 | 0,0000 |
| 28531 | DLSGQSSHPWNSNK | - | 48 | 0,0022 |
| 28534 | DLSGQSSHPWNSNK | - | 61 | 0,0001 |
| 28535 | DLSGQSSHPWNSNK | - | 49 | 0,0021 |
| 28570(D) | DLSGQSSHPWNSNK | - | 96 | 0,0000 |
| 28567(D) | DLSGQSSHPWNSNK | - | 73 | 0,0000 |
| 28571(D) | DLSGQSSHPWNSNK | - | 79 | 0,0000 |
| 28577(D) | DLSGQSSHPWNSNK | - | 48 | 0,0022 |
| 28574(D) | DLSGQSSHPWNSNK | - | 60 | 0,0002 |
| 28573(D) | DLSGQSSHPWNSNK | - | 55 | 0,0005 |
| 28576(D) | DLSGQSSHPWNSNK | - | 58 | 0,0003 |
| 28572(D) | DLSGQSSHPWNSNK | - | 94 | 0,0000 |
| 30906 | 'DLSGQSSHPWNSNK | - | 51 | 0,0000 |
| 30968(D) | 'DLSGQSSHPWNSNK | - | 31 | 0,0009 |
| 30973(D) | 'DLSGQSSHPWNSNK | - | 37 | 0,0003 |
| | PYDVPDYAPSSPSPSSD** | | | |

**b**

| Query | AUG-78/32-117CAG | | Ions Score | Expect |
|---|---|---|---|---|
| | **LVNVRYKSYFRRASEETRSLL*KTAAKAATAAAAAAAAAAAAAAAAA | | | |
| | AAAAAAAAAAAAAAAAAAAAAAAAAAAAAAAAAAAAAAAAAAAAAAAAAAAAAAAAAAAA | | | |
| 26136 | LISEE- | - | 99 | 0,0000 |
| 26134 | LISEE- | - | 49 | 0,0019 |
| 26137 | LISEE- | - | 58 | 0,0003 |
| 26138 | LISEE- | - | 90 | 0,0000 |
| 26139 | LISEE- | - | 74 | 0,0000 |
| 26140 | LISEE- | - | 82 | 0,0000 |
| 26142 | LISEE- | - | 42 | 0,0082 |
| 31016 | TEFTSMEFEQK | - | 48 | 0,0017 |
| 31768(O) | TEFTSMEFEQK | - | 57 | 0,0003 |
| 84318(O) | HAAAAAAAGPIRTEFTSMEFEQK | - | 35 | 0,0003 |
| | AAAAAAAAAAAAAAAAAAAAAAAAAAAAAAAAAAAAAAAAGPIRTEFTSMEFEQKLISEE | | | |
| 22885 | KLISEE- | - | | |
| 26136 | -DLLPIR | - | 99 | 0,0000 |
| 26134 | -DLLPIR | - | 49 | 0,0019 |
| 26137 | -DLLPIR | - | 58 | 0,0003 |
| 26138 | -DLLPIR | - | 90 | 0,0000 |
| 26139 | -DLLPIR | - | 74 | 0,0000 |
| 26140 | -DLLPIR | - | 82 | 0,0000 |
| 26142 | -DLLPIR | - | 42 | 0,0082 |
| | DLLPIRCSRLRSIITITI** | | | |
| 22885 | -DLLPI | | | |

**Fig. S8** Peptides identified by LC-MS/MS analysis from AUG-initiated polyQ and RAN polyA proteins expressed in cells transfected with AUG-78/32-117CAG. **a** Predicted amino acid sequence of canonically translated polyQ protein from AUG-78/32-117CAG construct is highlighted in orange. Peptides that are result of LysC digestions are shown in light blue, whereas peptides that are result of trypsin digestions are in black. **b** Predicted amino acid sequence of RAN polyA protein from AUG-78/32-117CAG construct is highlighted in orange. Peptides that are result of trypsin digestions are shown in light blue, whereas peptides that are result of elastase digestions are in black. Ion scores and the expected values corresponding to the identified peptides are indicated. Peptide amino acids sequences with their respective query identification numbers are shown, together with identified modifications i.e. A – N-terminal acetylation; D – Deamidation of Asn (N) and Gln (Q); O – oxidation of Met (M); G – Gln->pyro-Glu conversion. Modified amino acids are underlined, excluding N-terminal acetylation that is indicated by a red asterisk. Any amino acid predicted by LC-MS/MS that differs from the query amino acid sequence is highlighted in red. ‘ – isobaric glycine addition error as a result of N-terminal carbamidomethylation. Stop codons are indicated by black asterisks

### Fig. S9

**a**

| Query | 78/32-115CAG | | Ions Score | Expect |
|---|---|---|---|---|
| 3644 | REAYFEK | - | 51 | 0,0000 |
| 8195 | RREAYFEK | - | 34 | 0,0093 |
| 10536 | 'RREAYFEK | - | 35 | 0,0003 |
| 1172(A) | *SEELRK | - | 40 | 0,0001 |
| 4657 | LTSEELRK | - | 35 | 0,0004 |
| 2874(A) | *TSEELRK | - | 39 | 0,0001 |
| 11942 | TNLTSEELRK | - | 41 | 0,0001 |
| 11941 | TNLTSEELRK | - | 47 | 0,0000 |
| | **SGTNLTSEELRKRREAYFEKQQQKQQQQQQQQQQQQQQQQQQQQQQQ | | | |
| 16277(A) | *GTNLTSEELRK | - | 52 | 0,0000 |
| 5088 | LTSEELRK | - | 41 | 0,0001 |
| 2213 | LTSEELR | - | 41 | 0,0001 |
| 3293(A) | *TSEELRK | - | 34 | 0,0006 |
| 1262(A) | *SEELRK | - | 22 | 0,0067 |
| 983 | EAYFEK | - | 44 | 0,0011 |
| | QQQQQQQQQQQQQQQQQQQQQQQQQQQQQQQQQQQQQQQQQQQQQQQQQQQQQQQQQQQQ | | | |
| 27315 | DLSGQSSHPWNSNK | - | 43 | 0,0001 |
| 33931(D) | RDLSGQSSHPWNSNK | - | 32 | 0,0007 |
| 33929(D) | RDLSGQSSHPWNSNK | - | 48 | 0,0000 |
| 65725(DD) | SQQQQQQRDLSGQSSHPWNSNK | - | 33 | 0,0005 |
| | QQQQQQQQQQQQQQQQQQQQQQQQQQQQQQQQQQQQRDLSGQSSHPWNSNKNSSQKRICY | | | |
| 50816(D) | DLSGQSSHPWNSNKNSSQK | - | 35 | 0,0480 |
| 28145 | DLSGQSSHPWNSNK | - | 53 | 0,0008 |
| 28171(D) | DLSGQSSHPWNSNK | - | 99 | 0,0000 |
| 28173(D) | DLSGQSSHPWNSNK | - | 65 | 0,0000 |
| 30524 | 'DLSGQSSHPWNSNK | - | 49 | 0,0000 |
| | PYDVPDYAPSSPSPSSD** | | | |

**b**

| Query | 78/32-115CAG | | Ions Score | Expect |
|---|---|---|---|---|
| | **LVIRYKSYFRRASEETRSLL*KTAAKAATAAAAAAAAAAAAAAAAA | | | |
| | AAAAAAAAAAAAAAAAAAAAAAAAAAAAAAAAAAAAAAAAAAAAAAAAAAAAAAAAAAAA | | | |
| 28223 | LISEE- | - | 48 | 0,0020 |
| 28222 | LISEE- | - | 95 | 0,0000 |
| 28220 | LISEE- | - | 92 | 0,0000 |
| 28219 | LISEE- | - | 99 | 0,0000 |
| 33310 | TEFTSMEFEQK | - | 52 | 0,0008 |
| 34146(O) | TEFTSMEFEQK | - | 83 | 0,0000 |
| 90729 | NAAAAAAAAAAAAAAAAAAAAAAAAAAAAAGPIR | - | 55 | 0,0000 |
| | AAAAAAAAAAAAAAAAAAAAAAAAAAAAAAAAAAAAAAAAGPIRTEFTSMEFEQKLISEE | | | |
| 21187 | KLISEE- | - | ??? | ??? |
| 28223 | -DLLPIR | - | 48 | 0,0020 |
| 28222 | -DLLPIR | - | 95 | 0,0000 |
| 28220 | -DLLPIR | - | 92 | 0,0000 |
| 28219 | -DLLPIR | - | 99 | 0,0000 |
| | DLLPIRCSRLRSIITITI** | | | |
| 21187 | -DLLPI | | ??? | ??? |

**Fig. S9** Peptides identified by LC-MS/MS analysis from RAN polyQ and RAN polyA proteins expressed in cells transfected with 78/32-115CAG construct. **a** Predicted amino acid sequence of RAN polyQ protein from 78/32-115CAG construct is highlighted in orange. Peptides that are result of LysC digestions are shown in light blue, whereas peptides that are result of trypsin digestions are in black. **b** Predicted amino acid sequence of RAN polyA protein from 78/32-115CAG construct is highlighted in orange. Peptides that are result of trypsin digestions are shown in light blue, whereas peptides that are result of elastase digestions are in black. Ion scores and the expected values corresponding to the identified peptides are indicated. Peptide amino acids sequences with their respective query identification numbers are shown, together with identified modifications i.e. A – N-terminal acetylation; D – Deamidation of Asn (N) and Gln (Q); O – oxidation of Met (M); G – Gln->pyro-Glu conversion. Modified amino acids are underlined, excluding N-terminal acetylation that is indicated by a red asterisk. Any amino acid predicted by LC-MS/MS that differs from the query amino acid sequence is highlighted in red. ' – isobaric glycine addition error as a result of N-terminal carbamidomethylation. Stop codons are indicated by black asterisks

### Fig. S10

**a**

| Query | 33/32-119CAG | | Ions Score | Expect |
|---|---|---|---|---|
| | **AYFEKQQQKQQQQQQQQQQQQQQQQQQQQQQQ | | | |
| | QQQQQQQQQQQQQQQQQQQQQQQQQQQQQQQQQQQQQQQQQQQQQQQQQQQQQQQQQQQQ | | | |
| 22116 | LSGQSSHPWNSNK | - | 61 | 0,0000 |
| 22114 | LSGQSSHPWNSNK | - | 56 | 0,0000 |
| 27542 | DLSGQSSHPWNSNK | - | 55 | 0,0000 |
| 27541 | DLSGQSSHPWNSNK | - | 60 | 0,0000 |
| 34225(D) | RDLSGQSSHPWNSNK | - | 36 | 0,0003 |
| 34226(D) | RDLSGQSSHPWNSNK | - | 43 | 0,0001 |
| 34221(D) | RDLSGQSSHPWNSNK | - | 74 | 0,0000 |
| 34179 | RDLSGQSSHPWNSNK | - | 58 | 0,0000 |
| 34180 | RDLSGQSSHPWNSNK | - | 44 | 0,0001 |
| 34178 | RDLSGQSSHPWNSNK | - | 84 | 0,0000 |
| 36567(D) | 'RDLSGQSSHPWNSNK | - | 46 | 0,0000 |
| 36533 | 'RDLSGQSSHPWNSNK | - | 45 | 0,0000 |
| 44472 | QQRDLSGQSSHPWNSNK | - | 33 | 0,0005 |
| | QQQQQQQQQQQQQQQQQQQQQQQQQQQQQQQQQQQQRDLSGQSSHPWNSNKNSSQKRICY | | | |
| 72863(AD) | *LQQQQQQQRDLSGQSSHPWNSNK | - | 32 | 0,0010 |
| 76321(A/8xD) | *TQQQQQQQQRDLSGQSSHPWNSNK | - | 43 | 0,0001 |
| 68264(8xD) | NQQQQQQRDLSGQSSHPWNSNK | - | 38 | 0,0002 |
| 61818(O/DDD) | MQQQQRDLSGQSSHPWNSNK | - | 32 | 0,0018 |
| 51408(D) | DLSGQSSHPWNSNKNSSQK | - | 47 | 0,0031 |
| 28622 | DLSGQSSHPWNSNK | - | 94 | 0,0000 |
| 28624 | DLSGQSSHPWNSNK | - | 64 | 0,0001 |
| 28626 | DLSGQSSHPWNSNK | - | 61 | 0,0001 |
| 28627 | DLSGQSSHPWNSNK | - | 53 | 0,0008 |
| 28623 | DLSGQSSHPWNSNK | - | 35 | 0,0460 |
| 28660(D) | DLSGQSSHPWNSNK | - | 107 | 0,0000 |
| 28661(D) | DLSGQSSHPWNSNK | - | 69 | 0,0000 |
| 28664(D) | DLSGQSSHPWNSNK | - | 61 | 0,0001 |
| 28662(D) | DLSGQSSHPWNSNK | - | 46 | 0,0042 |
| 28663(D) | DLSGQSSHPWNSNK | - | 52 | 0,0009 |
| 28693(DD) | DLSGQSSHPWNSNK | - | 47 | 0,0033 |
| 40886 | MRDLSGQSSHPWNSNK | - | 40 | 0,0002 |
| 31076 | 'DLSGQSSHPWNSNK | - | 97 | 0,0000 |
| 31081 | 'DLSGQSSHPWNSNK | - | 41 | 0,0001 |
| 31137(D) | 'DLSGQSSHPWNSNK | - | 100 | 0,0000 |
| | PYDVPDYAPSSPSPSSD** | | | |

**b**

| Query | 33/32-119CAG | | Ions Score | Expect |
|---|---|---|---|---|
| | **LVSLL*KTAAKAATAAAAAAAAAAAAAAAAA | | | |
| | AAAAAAAAAAAAAAAAAAAAAAAAAAAAAAAAAAAAAAAAAAAAAAAAAAAAAAAAAAAA | | | |
| 26873 | LISEE- | - | 47 | 0,00320 |
| 26872 | LISEE- | - | 95 | 0,00000 |
| 26870 | LISEE- | - | 64 | 0,00005 |
| 26869 | LISEE- | - | 96 | 0,00000 |
| 26868 | LISEE- | - | 68 | 0,00002 |
| 26867 | LISEE- | - | 38 | 0,02200 |
| 26871 | LISEE- | - | 99 | 0,00000 |
| 82620(OO) | MTSMEFEQKLISEE- | - | 51 | 0,00001 |
| 32576(O) | TEFTSMEFEQK | - | 63 | 0,00007 |
| | AAAAAAAAAAAAAAAAAAAAAAAAAAAAAAAAAAAAAAAAGPIRTEFTSMEFEQKLISEE | | | |
| 21117 | KLISEE- | - | ??? | ??? |
| 5907 | SEE- | - | ??? | ??? |
| 949 | SIITITI | - | 39 | 0,02000 |
| 26873 | -DLLPIR | - | 47 | 0,00320 |
| 26872 | -DLLPIR | - | 95 | 0,00000 |
| 26870 | -DLLPIR | - | 64 | 0,00005 |
| 26869 | -DLLPIR | - | 96 | 0,00000 |
| 26868 | -DLLPIR | - | 68 | 0,00002 |
| 26867 | -DLLPIR | - | 38 | 0,02200 |
| 26871 | -DLLPIR | - | 99 | 0,00000 |
| 82620 | -DLLPIR | - | 51 | 0,00001 |
| | DLLPIRCSRLRSIITITI** | | | |
| 21117 | -DLLPI | - | ??? | ??? |
| 5907 | -DLLPI | - | ??? | ??? |

**Fig. S10** Peptides identified by LC-MS/MS analysis from RAN polyQ and RAN polyA proteins expressed in cells transfected with 33/32-119CAG construct. **a** Predicted amino acid sequence of RAN polyQ protein from 33/32-119CAG construct is highlighted in orange. Peptides that are result of LysC digestions are shown in light blue, whereas peptides that are result of trypsin digestions are in black. **b** Predicted amino acid sequence of RAN polyA protein from 33/32-119CAG construct is highlighted in orange. Peptides that are result of trypsin digestions are shown in light blue, whereas peptides that are result of elastase digestions are in black. Ion scores and the expected values corresponding to the identified peptides are indicated. Peptide amino acids sequences with their respective query identification numbers are shown, together with identified modifications i.e. A – N-terminal acetylation; D – Deamidation of Asn (N) and Gln (Q); O – oxidation of Met (M); G – Gln->pyro-Glu conversion. Modified amino acids are underlined, excluding N-terminal acetylation that is indicated by a red asterisk. Any amino acid predicted by LC-MS/MS that differs from the query amino acid sequence is highlighted in red. ‘ – isobaric glycine addition error as a result of N-terminal carbamidomethylation. Stop codons are indicated by black asterisks

#### Fig. S11

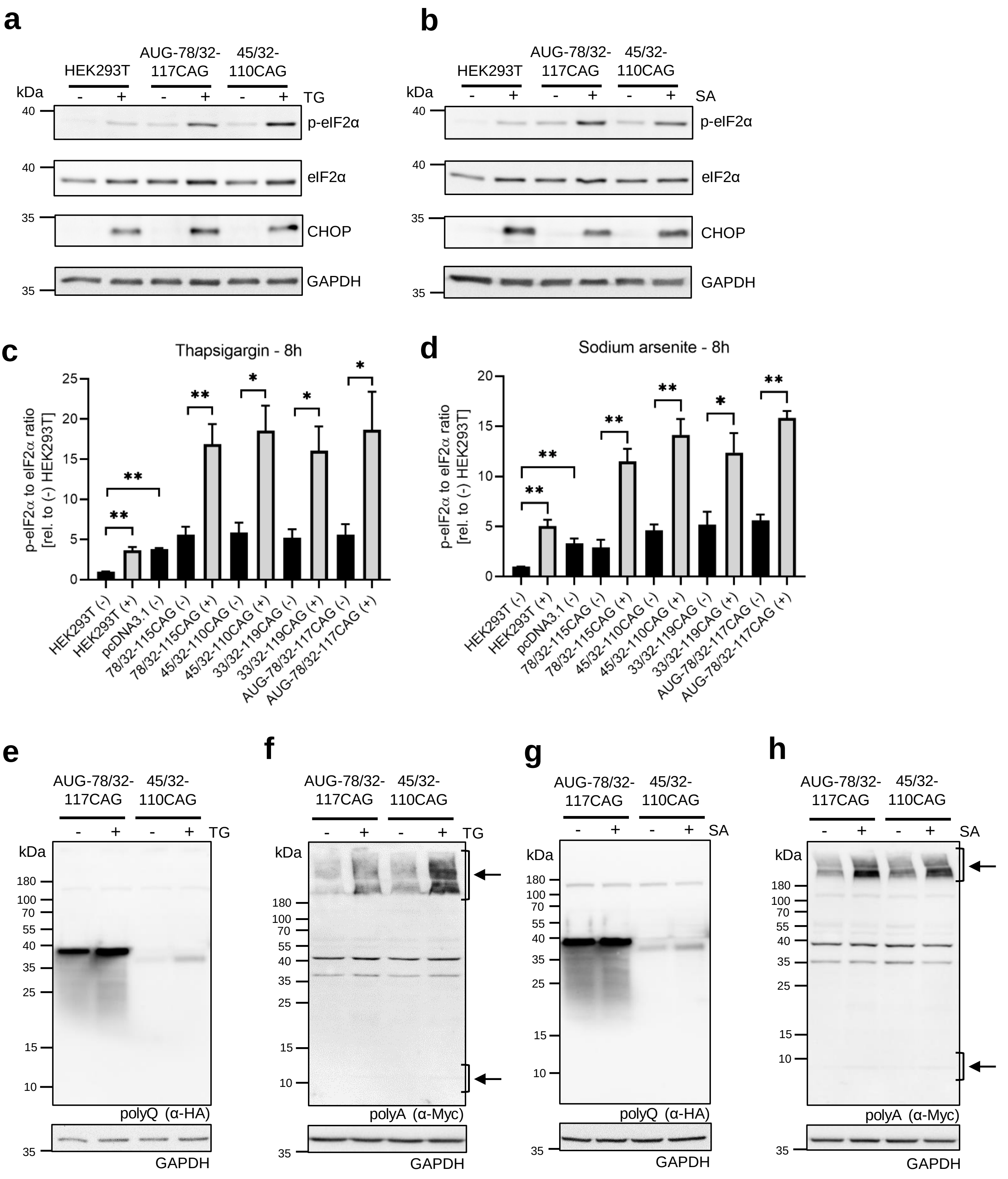
a
HEK293T
AUG-78/32-117CAG
45/32-110CAG
kDa
TG
p-eIF2α
eIF2α
CHOP
GAPDH
b
SA
c
Thapsigargin - 8h
p-eIF2α to eIF2α ratio [rel. to (-) HEK293T]
HEK293T (-)
HEK293T (+)
pcDNA3.1 (-)
78/32-115CAG (-)
78/32-115CAG (+)
45/32-110CAG (-)
45/32-110CAG (+)
33/32-119CAG (-)
33/32-119CAG (+)
AUG-78/32-117CAG (-)
AUG-78/32-117CAG (+)
d
Sodium arsenite - 8h
e
f
g
h
polyQ (α-HA)
polyA (α-Myc)

**Fig. S11** Canonical and SCA3 RAN translation upon cellular stress induction. For all the analyses presented in this figure, lysates were obtained from HEK293T cells 20 h after their transfection with the indicated constructs followed by 8 h of treatment with 1 μM thapsigargin (TG) or 50 μM sodium arsenite (SA). **a, b** Western blot analysis of the phospho-eIF2α at the Ser51 site, eIF2α and CHOP in the HEK293T cells that were either transfected or not with the indicated constructs and treated or not with TG or SA. **c, d** The ratio of phospho-eIF2α to total eIF2α after treatment or not with TG or SA and normalized to the ratio of the phospho-eIF2α to total eIF2α of the untransfected and untreated HEK293T cells. The graph bars represent the mean value ± SEM from 5 biological replicates. Two-tailed *t* test $*p<0.05$, $**p<0.005$. **e, f** Expression of the AUG-initiated polyQ, RAN polyQ and RAN polyA proteins after treatment or not with TG was analyzed by western blotting using anti-HA and anti-c-Myc antibodies, respectively. **g, h** Expression of the AUG-initiated polyQ, RAN polyQ and RAN polyA proteins after treatment or not with SA was analyzed by western blotting using anti-HA and anti-Myc antibodies, respectively. Specific RAN polyA products used in the quantitative analysis are indicated by black arrows. For all the western blot analyses, GAPDH was used a loading control
